# Gibbs Process Determines Survival and Reveals Contact-Inhibition Genes in Glioblastoma Multiforme

**DOI:** 10.1101/608414

**Authors:** Lavanya Kannan, Tarjani Agarwal, Matija Snuderl, David Zagzag, Erik Sulman, Jason Huse, Kasthuri Kannan

## Abstract

Tumor growth is a spatiotemporal birth-and-death process with loss of heterotypic contact-inhibition of locomotion (CIL) of tumor cells promoting invasion and metastasis. Therefore, representing tumor cells as two-dimensional points, we can expect the tumor tissues in histology slides to reflect realizations of spatial birth-and-death process which can be mathematically modeled to reveal molecular mechanisms of CIL, provided the mathematics models the inhibitory interactions. Gibbs process as an inhibitory point process is a natural choice since it is an equilibrium process of the spatial birth-and-death process. That is if the tumor cells maintain homotypic contact inhibition, the spatial distributions of tumor cells will result in Gibbs hard core process over long time scales. In order to verify if this is the case, we applied the Gibbs process to 411 TCGA Glioblastoma multiforme patient images. Our imaging dataset included all cases for which diagnostic slide images were available.

The model revealed two clusters, one of which - the “Gibbs cluster,” showed the convergence of the Gibbs process with significant survival difference. Further smoothing the discretized (and noisy) inhibition metric, for both increasing and randomized survival time, we found a significant association of the patients in the Gibbs cluster with increasing survival time. The mean inhibition metric also revealed the point at which the homotypic CIL establishes in tumor cells. Besides, RNAseq analysis between patients with loss of heterotypic CIL and intact homotypic CIL in the Gibbs cluster unveiled cell movement gene signatures and differences in Actin cytoskeleton and RhoA signaling pathways as key molecular alterations. These genes and pathways have established roles in CIL. Taken together, our integrated analysis of patient images and RNAseq data provides for the first time a mathematical basis for CIL in tumors, explains survival as well as uncovers the underlying molecular landscape for this key tumor invasion and metastatic phenomenon.

## Introduction and preliminaries

Contact-inhibition of locomotion (CIL) is a fundamental mechanism that immobilizes cells in healthy tissues [3] and has long been implicated in the invasion of cancer cells [29]. When collisions happen between the same cell type, this mechanism is referred to as *homotypic* CIL, and *heterotypic* CIL occurs when the collisions are between different cell types. Several studies have demonstrated a loss of heterotypic CIL between tumor and normal cells and have hypothesized this as a phenomenon behind tumor invasion and metastasis [19] [4] [25]. At the same time, it is known that homotypic CIL is maintained between the tumor cells [18] [20]. Hence, we can expect the spatial distribution of cells in tumor tissue to reflect one or both of these phenotypes that would determine the survival of the patients, with loss of heterotypic CIL patients evidencing low survival due to tumor invasion and metastasis.

Although CIL is a dynamic mechanism that happens in real-time, by examining the spatial distribution of cells in tumor tissues, we can infer homotypic CIL and/or loss of heterotypic CIL. After all, if homotypic inhibition is maintained, the tumor cells should not be too clustered. Similarly, if the loss of heterotypic inhibition occurs, we can expect a clustering phenotype. Even though one cannot infer the mechanisms of homotypic or loss of heterotypic CIL based on dispersion or clustering phenotypes alone, we can deduce these processes if cell movement gene expression is also altered. Therefore, by examining the spatial statistics and applying appropriate inhibitory or cluster models, homotypic CIL or loss of heterotypic CIL can be inferred by integrating pathology images with gene expression data. Even though CIL as a collective migration of tissues has been a subject of extensive mathematical modeling [37], application of the stochastic process to understand these mechanisms in tumors through radiomics approaches has not been undertaken, and the mathematical description of CIL as a spatial point process remains elusive.

From a mathematical standpoint, several stochastic models for tumor evolution has been proposed since 1950’s. Five dominant paradigms have emerged [24], out of which Two-Stage Clonal Expansion (TSCE) and Moran models have been predominantly applied to study tumor growth [6], [22], [23]. TSCE posits that tumor growth follows two stages, an initiating stage which is modeled as an inhomogeneous Poisson process (because tumor initiation is a rare event) and proliferation/malignancy stage that follows the birth-and-death process. Thus, the Moran model also being a birth-and-death process, allows us to infer tumor growth is a birth-and-death process with CIL as a critical mechanism for invasion and metastasis.

Therefore, representing tumor cells as two-dimensional points, we can expect the tumor tissues in histology slides to reflect realizations of spatial birth-and-death process where each point/cell would maintain a certain distance from its neighbors if homotypic CIL is maintained. These spatial points can be mathematically modeled using an appropriate spatial point process that considers inhibitory interactions, and *Gibbs hard core process* as an inhibitory process to which birth-and-death process converges is a natural choice. A thorough treatment of spatial point processes and in particular Gibbs process is mathematically sophisticated, and beyond the scope of this article, however, we will illustrate the central ideas behind Gibbs process and related statistics transliterated in the language of cells and CIL. These are adapted (sometimes ad verbatim) from excellent introductory point process resources such as [1] and transliterated for our purposes.

### Overview of the Gibbs process and L-function (transliterated to reflect biological context)

We will use the word *intensity* to refer to the number of cells per unit area in a randomly chosen small region of a histology slide.

#### Gibbs hard core process

A spatial distribution of a finite collection of cells with intensity *β* can be assigned a probability density with respect to a completely random collection of cells with unit intensity. This probability density will change depending on the interaction between the cells. In the case of intact homotypic CIL in tumors, the cells are forbidden to come too close to each other, and when they maintain a mean distance *h*, the locations of the centers of the cells form a point process in which each pair of points is approximate *h* units apart. Such inhibition is called a *hard core* constraint, and the Gibbs hard core process is defined by the probability density when it satisfies the hard core constraint.

It is noteworthy to mention that there is a deep relationship between the spatial birth-and-death process and the Gibbs hard core process. Preston [27] proves, under the condition that there is a certain distance maintained between the cells, the spatial birth-and-death process will converge to a realization of the Gibbs hard core process. In particular, we have the following result:

Suppose that, in each small interval of time Δ*t*, each existing cell has probability *d*(*t*)Δ*t* of undergoing cell death, where *d*(*t*) is the apoptosis rate per cell per unit time. In the same time interval, in any small region of area Δ*a*, let a cell division take place with probability *m*(*t*)Δ*a*Δ*t*, where *m*(*t*) is the mitotic rate of the cell per unit time. Provided the dividing cell lies *h* units away from its nearest neighbor, no matter what the initial state of the cells are, over long time scales, this spatial birth-and-death process would reach an equilibrium in which any snapshot of the cells will be a realization of the Gibbs hard core process with Gibbs process intensity *β* = *m*(*t*)/*d*(*t*) and hard-core constraint *h*.

We can now see that when homotypic CIL is maintained in tumors, the hard core constraint is naturally satisfied and therefore if we represent the cells as two-dimensional points (segmented from histology slides), the Gibbs process is instinctive to apply. However, we do not know apriori if homotypic CIL is maintained for all tumors and we can expect some of the tumors to exhibit loss of heterotypic CIL (or both). The validity of the Gibbs model as an inhibitory process can be applied to test and distinguish the maintenance of homotypic CIL versus the loss of heterotypic CIL. Moreover, the validity of the model can be determined by the statistical theory that suggests the inference for the Gibbs models should be based on summary statistics such as the *L*-function [1].

#### L-function

The *L*-function is the commonly used transformation of the *K*-function. *K*-function is one of the very popular statistics for analyzing the spatial correlations between point patterns [1] proposed by Ripley [28]. The *K*-function, *K_r_*, of a randomly chosen small region of a histology slide image is the expected number of neighbors within radius *r* of a typical cell at location *u*, divided by the intensity *β*. Note that if we do not divide by the intensity of *β*, the expected number of neighbors within radius *r* will vary depending on the scale of the region. That is, if the slides are zoomed out/in, the expected number of *r*-neighbors will change. Dividing by the intensity *β* assures us the *K*-function is independent of scaling operations.

Mathematically,

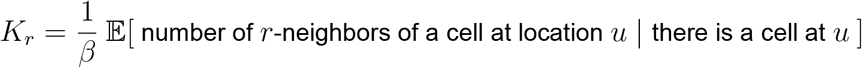

for any *r* ≥ 0 and any location *u*. The theoritical *K*-function of a homogeneous distribution of cells (aka homogeneous Poisson process) is *πr*^2^ and this quantity can be used to determine if the cells are clustered, or inhibited. For such a comparison, the transformed *K*-function, namely the *L*-function,

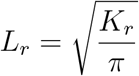

can be tested against the *L*-function of the homogeneous Poisson process, namely, 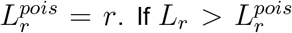 the cells would exhibit a clustering phenotype and 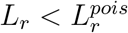, the cells will be contact-inhibited.

As noted above, if the cells are contact-inhibited, the validity of the Gibbs process can be determined by the summary statistics, the *L*-function. That is, if the cells are contact-inhibited, we can expect the fitted optimal value of Gibbs hard-core constraint, opt_*r*_ *G_r_*, to be approximately the same as the optimal value of | *L_r_* − *r* |, indicating the model fits the statistics as determined by the *L*-function. Therefore, we will use these metric for our results, that is *L_r_* ≡ opt_*r*_ | *L_r_* − *r* | and *G_r_* ≡ opt_*r*_*G_r_*. Thus, it is natural to study the metric *M*(*r*) = (*L_r_*, *G_r_*) to determine the convergence of the Gibbs process. We will refer *M*(*r*) as the *inhibition metric*. Further, for our analysis, we will use the smoothed one-dimensional version of the metric, which we call the *mean inhibition metric*, *MIM*(*r*), defined as:

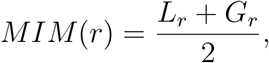

which we will use to analyze overall survival of the patients.

## Image processing

Figure 1 shows the image processing protocol employed to extract the point pattern to fit the Gibbs process for the TCGA images (image TCGA-02-0339 shown for illustration). We processed all images for which diagnostic slides were available (389 images, see ‘query.png’ in Supplemental Information) and 22 cases were added to further test the association of MGMT-promotor methylation found to be associated with one of the clusters (eventually a false association) in the initial 389 cohort, resulting in 411 patient images. Three slides, namely, TCGA-19-1388, TCGA-19-0963 and TCGA-19-1389 had bad diagnostics images, and so we used tissue slides. Each image was resolved until 50 *μm* and a region of 300 *μm*^2^ was randomly chosen and extracted for segmenting the cells.

**Figure 1:**
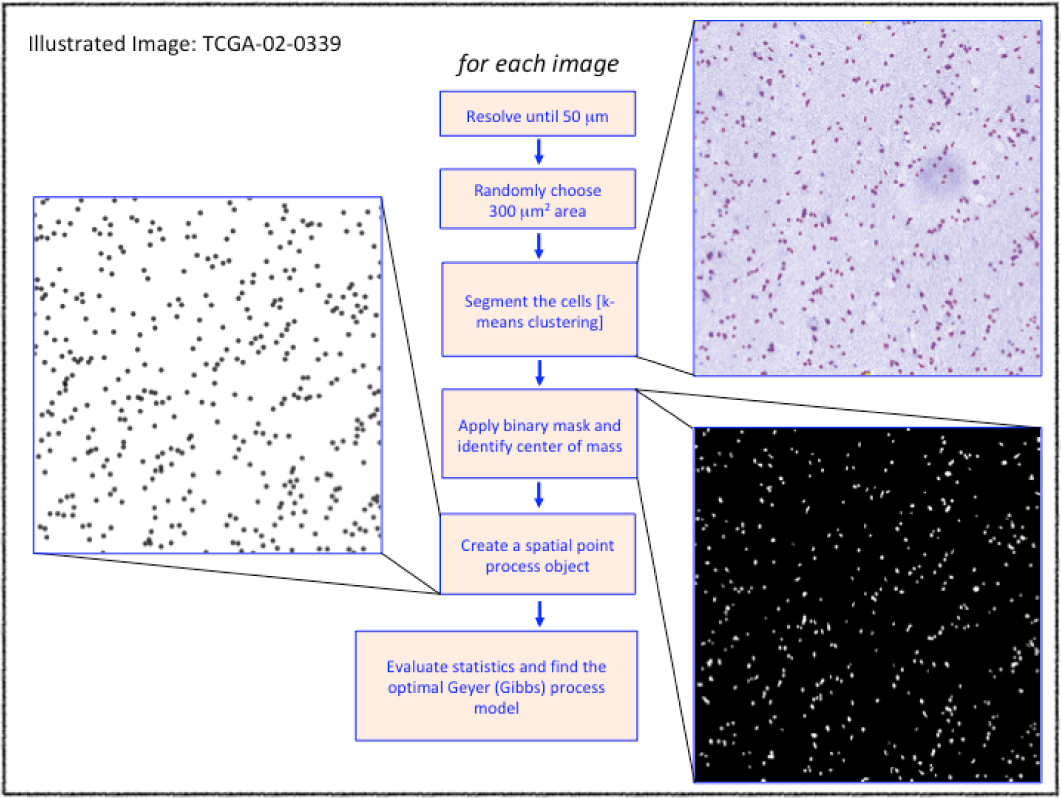
Image processing protocol

We note that the downstream analysis is independent of the resolution and the area since Ripley’s *K*-function (and hence the estimated *L*-function) gives a scale-free description of the expected number of cells within a neighborhood of an arbitrary cell in any region of interest [16]. The cells in the isolated regions were segmented using *k*-means segmentation algorithm; the center of masses was identified in the binary mask to generate a spatial point pattern corresponding to the images. A generalized form of the Gibbs hard core process, namely the Geyer saturation process, was fitted to the point pattern for downstream analysis (refer to ‘Gibbs_Models.pdf’ document in Supplemental Information for the type of Gibbs model that was applied). The model parameters and the L-function statistics are presented in Supplemental Data 01.

## Results

Here we discuss the results after we obtained the *G_r_* and *L_r_* values by processing all the images using the image processing protocol described above. Both *L_r_* and *G_r_* values were scaled using the standardized *z*-score for the rest of the analysis.

### Convergence of the Gibbs Process

As noted above, if the Gibbs model fits the data, we would expect the *L_r_* and *G_r_* values to be close and hence the metric *M*(*r*) = (*G_r_*, *L_r_*) is likely to cluster for some patients. We determined the optimal number of clusters for this metric. Figure 2 (a) shows the result of applying 30 indices for determining the optimal number of clusters (see Methods section for the algorithm used). A majority of the indices reported the optimal number of clusters as 2 for this metric. One of the clusters, we call Cluster 1, had a smaller range of values for this metric than the other cluster which we refer as the “Gibbs cluster.” Figure 2 (b) shows the metric *M*(*r*) for Cluster 1 (red) and the Gibbs cluster (blue) where 145 patients were in Cluster 1 and 266 belonged to the Gibbs cluster (see Supplemental Data 02 for cluster association). We note that the variance of *M*(*r*) in the Gibbs cluster is lesser than Cluster 1, indicating that *G_r_* and *L_r_* values converge at least for the Gibbs cluster. Therefore, in order to determine if this is a convergence behavior in mathematical terms, we plotted the *k*-means centroids as we included more *M*(*r*) values. Figure 2 (c) shows the *k*-means centroids starting with 20 patients and increasing to 411 patients (see Supplemental Data 04 for the centroids of 392 patients). We see the two subgroups of patients, one with “clustering” phenotype (Cluster 1) and the other “inhibitory” phenotype (Gibbs cluster). Moreover, calculating the overall survival time for these two clusters of patients from the TCGA database, revealed the patients with the clustering phenotype survive significantly shorter than the patients with inhibitory phenotype. This is illustrated in Figure 2 (d) (also see Supplemental Data 05 for the actual numbers). One can expect this result since the patients with loss of heterotypic CIL between tumor and normal cells that result in tumor cell invasion and metastasis are likely to have clustering phenotype and poor survival. On the other hand, patients with homotypic CIL between tumor cells, where cells repel each other, are less prone to tumor cell invasion leading to high survival.

**Figure 2:**
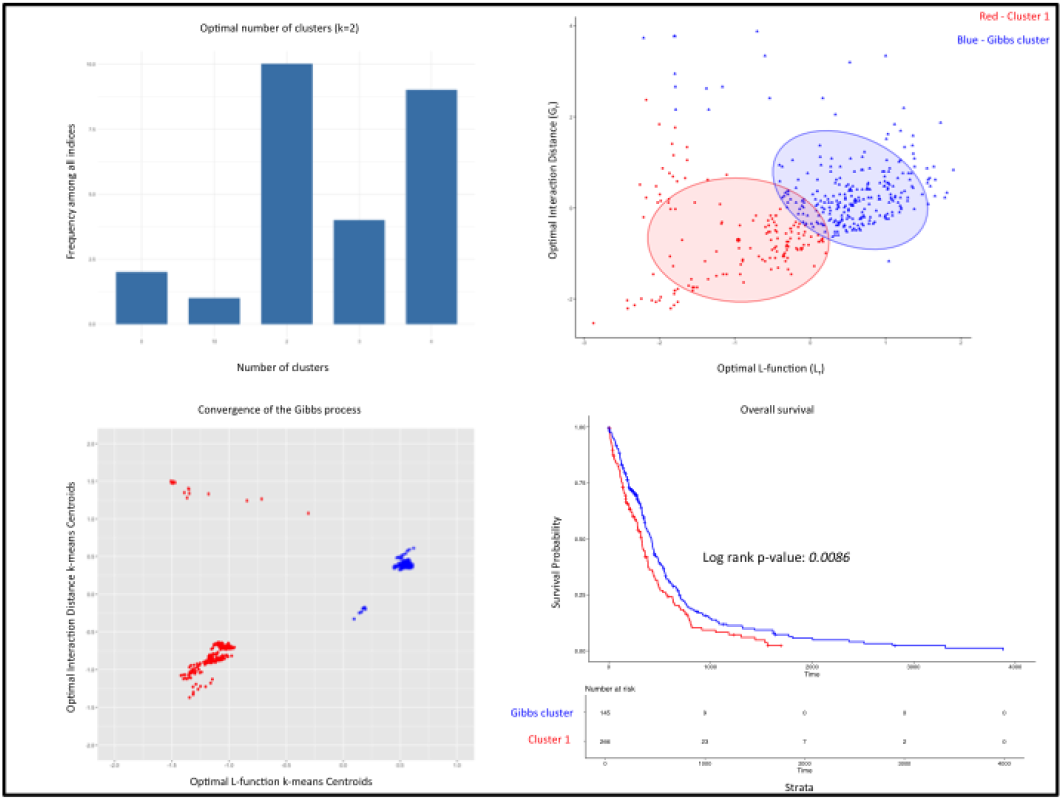
Convergence of Gibbs process

### Gibbs process determines survival time

Although the difference in the overall survival of the patients is significant between the two groups, we found several patients in the Gibbs cluster with poor survival and patients in Cluster 1 with high survival. This finding is contrary to the expectation that if the loss of heterotypic CIL and homotypic CIL distinguishes these patient types, theoretically, the number of patients in the Gibbs cluster who do worse should be very small and correspondingly, the number of patients in Cluster 1 who survive high should be negligible as well.

In other words, the log-rank statistical significance between the groups must be much higher than what it is currently presented. We hypothesized that this could be due to the noise in the metric *M*(*r*) and smoothing the metric according to increasing survival time in each cluster will likely delineate and identify the mark where actual homotypic CIL is established in patients. Figure 3 (a) shows the one dimensional version of the smoothed metric *M*(*r*), i.e., *MIM*(*r*), using the Loess regression fitting for the increasing survival time (dark blue), and randomized survival time (dashed-gray) patients, where increasing survival time and randomized survival time are determined in each cluster (the codes and the data for generating this are also provided). It also shows the randomized *k*-means centroids for the two clusters as horizontal dots, red and blue.

**Figure 3:**
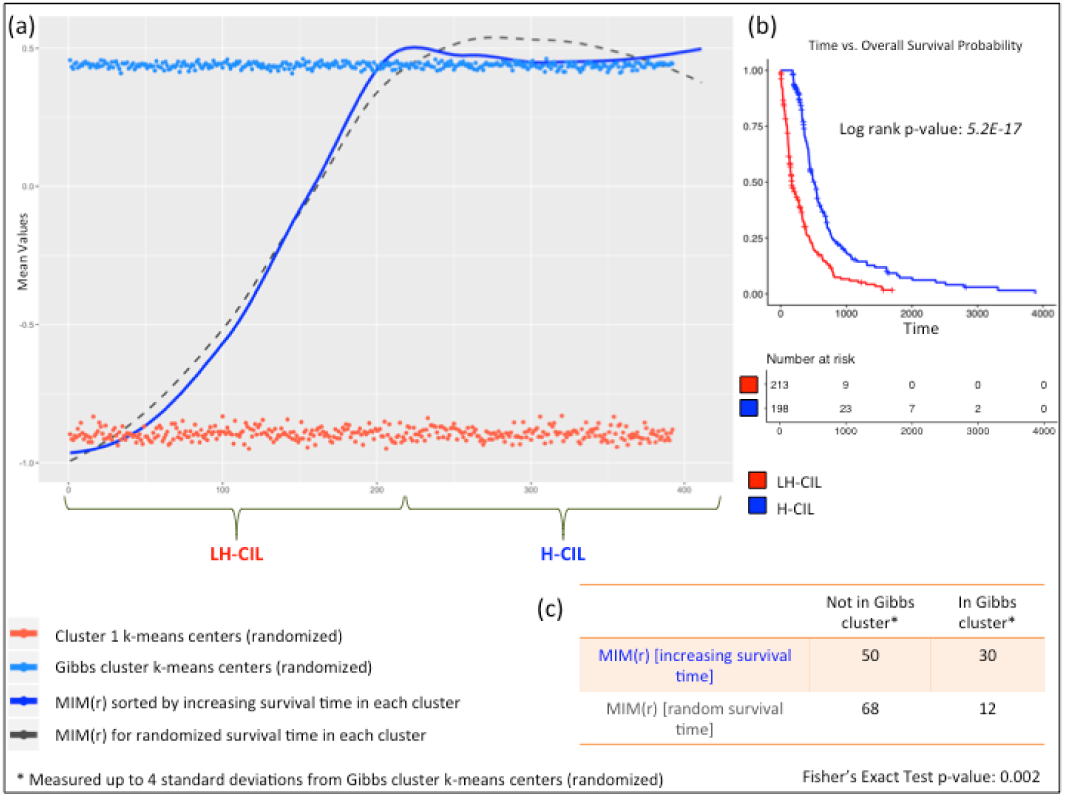
Survival time analysis

We note that *MIM*(*r*) with increasing survival time is significantly associated (Figure 3 (c), Fisher’s exact test 0.002) with the randomized *k*-means centroids of the Gibbs cluster than *MIM*(*r*) with randomized survival time, demonstrating that it is precisely the noise in the metric *M*(*r*) that included many patients with poor survival time in the Gibbs cluster and high survival time in Cluster 1. This association was measured up to 4 standard deviations from the randomized *k*-means centroids of the Gibbs cluster. Moreover, the association of *MIM*(*r*) with increasing survival time also revealed the point at which the homotypic CIL establishes in the tumor cells - around 179 days, corresponding to index 213 in the Gibbs cluster (see Supplemental Figure 01 and Supplemental Data 06). Using this time-point to distinguish loss of heterotypic CIL (LH-CIL) and homotypic CIL (H-CIL), we see that patients with LH-CIL have an extremely poor outcome than H-CIL patients as shown in Figure 3 (b).

### Gene expression signatures between loss of heterotypic CIL and homotypic CIL subgroups

In order to determine if there are expression differences between LH-CIL and H-CIL subgroups, we performed RNA-seq analysis from the TCGA data that was available in the GDC database for several of the patients. There were 22 patients in Cluster 1 and 47 patients in the Gibbs cluster who had the corresponding RNA-seq data (see Supplemental Data 07 and 08). The analysis revealed 447 differentially expressed molecules several of which were significantly implicated in cell movement, cell adhesion, and binding (see Supplemental Data 09). More specifically, 48 out of the 86 genes involved in cell movement were found to have low enrichment in H-CIL group (green) as compared to LH-CIL group (red) and exhibited decreased cell movement consistent with several findings [32][34][17][33] that show these genes increase cell migration and invasion (see Supplemental Figure 02, 03, 04 and Supplemental Data 10). This can be directly attributed to homotypic CIL that results in inhibition between the tumor cells. Figure 4 (a) shows the differential gene expression for cell movement genes with an absolute value of the average gene expression difference between LH-CIL and H-CIL groups greater than 0.2. Supplemental Figure 07 and Supplemental Data 11 shows the heatmap and gene expression values (original and transformed) for the signature derived in Figure 4 (a). Correspondingly, the increased expression of these 48 cell movement genes in LH-CIL patients demonstrate loss of heterotypic CIL leading to tumor invasion, and hence poor survival.

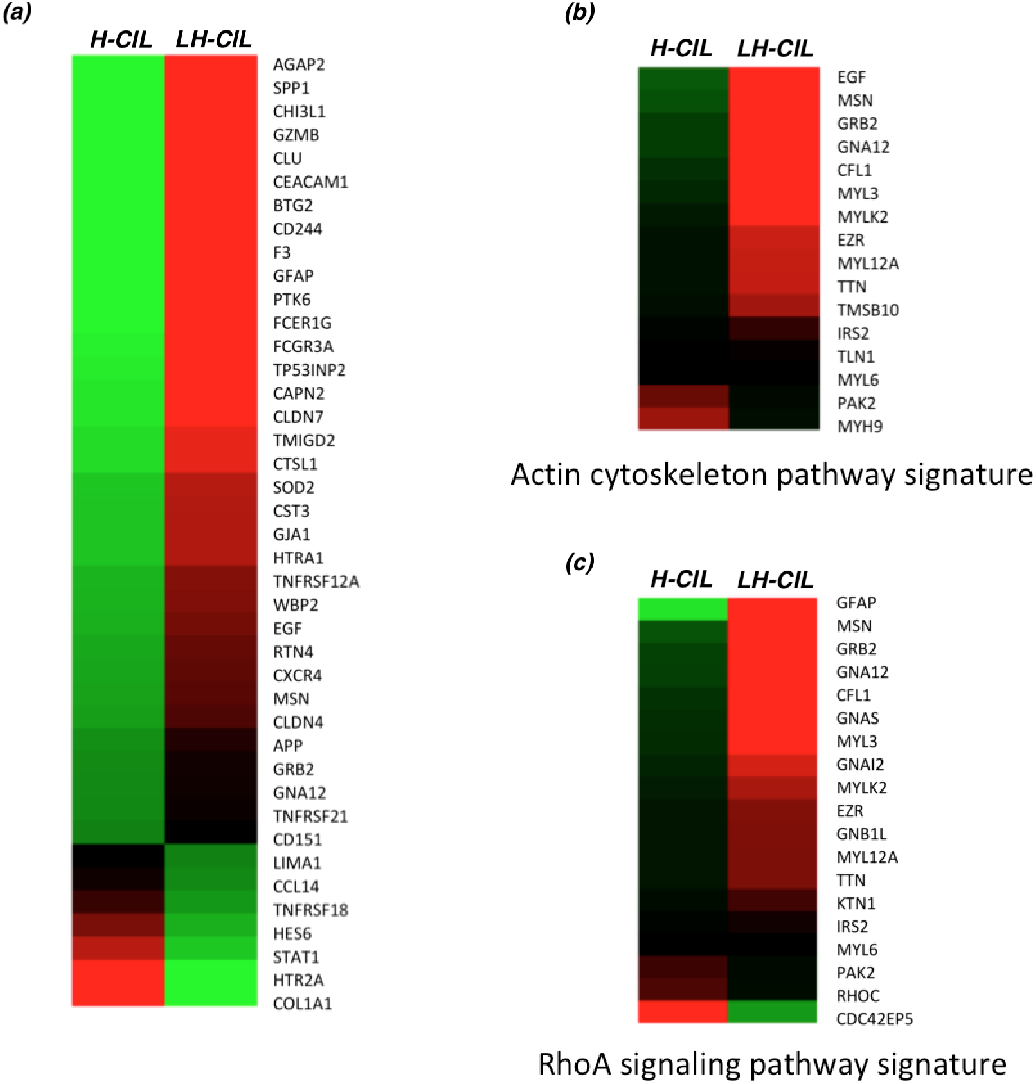

Further, pathway analysis using the differentially expressed genes revealed genes in Actin cytoskeleton and RhoA signaling as key molecular alterations (Figure 4 (b) and (c)). Supplemental Data 12-15 provides the values of these gene expression signatures and Supplemental Figure 05, and 06 shows the implicated signaling pathways. The actin cytoskeleton plays a critical role in cell movement as they link to extracellular matrix proteins (through adhesions) when cell protrusions stabilize during movement [10]. Also, it is known that the three Rho isoforms – RhoA, RhoB, and RhoC – can induce stress fiber formation [20] [35]. Since directional cell migration involves the polarized formation of cell protrusions in the front and contraction of stress fibers at the edge, our results suggest that unstable protrusions and/or deformed stress fibers in LH-CIL patients are likely to induce tumor cell invasion. On the other hand, over-stabilized protrusions and/or formation of additional stress fibers may result in an enhanced CIL in homotypic patients.

## Discussion

Spatial point processes are powerful mathematical frameworks for studying point patterns. Cells invariably move, and by representing them as points, it is appropriate to study these movement behaviors using point process. Gibbs process as an inhibition model is a natural choice for studying inhibitory interactions that drive CIL in tumors. By fitting the Gibbs process to diagnostic slide images of Glioblastoma multiforme, for the first time, we have introduced a mathematical framework for CIL in tumors and have validated the hypothesis presented in [20], to quote - “the invasive behavior of tumors is facilitated by the absence of heterotypic CIL with normal cells, whereas homotypic CIL between cancer cells helps collective migration and/or dispersion of the tumor.”

Even though CIL was first characterized through the work of Abercrombie and Heaysman six decades earlier [2], and has been widely studied model for cell migration and motility in developmental biology (especially in the development of neural crests), its critical role in cancers is yet to be understood. Although we know cancer is primarily a disease of cell invasion and CIL has been hypothesized as the principal mechanism for clonal expansion and metastasis, the molecular mechanisms of CIL have not been elucidated yet. The power of our approach is that these mechanisms can be determined only using microscopy images along with a robust mathematical model. In particular, the fact that tumor cells invade and this invasion can be deciphered through images presents a dominant paradigm for cancer diagnostics and treatment options. It opens a window for a plethora of investigations, both clinical and biological, that can be followed when integrated with the existing knowledge on the molecular landscape of Glioblastoma multiforme. Such integration can be made using the marked spatial point processes such as marked Gibbs or marked Geyer saturation process.

Although we demonstrated the effectiveness of this approach in Glioblastoma multiforme, it is important to note that the foundations of our work rely only on images and mathematical models and therefore can be applied to any cancer types. Gibbs and Strauss’s models are applicable where cell-to-cell ‘repulsion’ interactions define heterogeneity like in diffuse gliomas. In tumors of epithelial origin, tumor microenvironment support clustering of cells in a process called “nesting.” For these tumors or even if there is clustering in diffuse gliomas, probability models such as Cox, *Neyman-Scott, Matérn, Thomas, Gauss-Poisson, Cauchy* and several other processes can be applied and compared. Moreover, cluster point processes are natural to apply and compare in the setting of clonal evolution since these processes model parent-offspring associations that mimic the natural process of cell division.

It is intriguing that no such comparative analysis of different point process models have been performed purely on histology images (that is, even ignoring molecular data). This is not to say point process models have not been used in the study of tumors. Although there have been efforts to learn tumor’s spatial architecture purely from a stochastic process perspective and point processes has been applied in various cancer studies [14][13][30][7], fitting models to diagnostic images has not been attempted. Moreover, none of the studies have obtained molecular signatures through model fitting approach despite that there are several software packages, including *spatstat* [5], that are available for spatial point process modeling. As a comprehensive point process modeling package, *spatstat* also allows easy integration of genetic alterations as a marked point process enabling *multitype* comparative analysis, where we can analyze the effect of interactions between several mutation types. Processes such as *Multitype Gibbs, Multitype Strauss*, and *area-interaction* models allow higher-order interactions between several mutation types and these processes can be implemented to study heterogeneity and clonal evolution on the spatial level leading to translational medicine.

In short, spatial point process models establish the causal relationship between biology and contemporary radiomic approaches in medicine where the primary goal is computing the correlations between computational features and clinical outcomes. Without establishing a causal relationship, translating the results of radiomics into a clinical setting is likely to be unachievable [36], and by integrating spatial point process models, effective radiomic systems can be developed and implemented to translate biological findings to clinical care.

## Methods

### Diagnostic slide images

We queried the GDC portal (https://portal.gdc.cancer.gov/) for GBM cases for which diagnostic cases were available. The query is presented in Supplemental Information.

### Image and data processing

Microscopy H&E images were resolved until 50 *μm* and a region of 300 *μm*^2^ was randomly chosen using QuPath [26], an open source software for digital pathology image analysis. Image segmentation was done using color-based segmentation algorithm that uses *k*-means clustering in Matlab [31]. Spatial statistics such as the L-function and the parameters of the Geyer saturation process (a Gibbs model) were obtained using spatstat [1], an open source software for spatial statistics.

For determining the optimal number of clusters, we used NbClust [8] algorithm that applies 30 indices to estimate the optimum. The metrics for our data set are given in Supplemental Data 03. Also, for the *k*-means clustering, we used the standard package *k*-means as implemented in the R programming language.

### Gene expression and pathway analysis

RNA-seq analysis was performed using TCGAbiolinks [9] queries which facilitate the GDC open-access data retrieval to perform standard reproducible differential expression analysis. TCGAbiolinks uses EdgeR differential expression package [21] under the query *TCGAanalyze_DEA*. We used false discovery rate cutoff of 0.001 and log-fold change cutoff of 6 with the generalized linear model *glmLRT* as parameters to TCGAanalyze_DEA. This resulted in 447 differentially expressed genes (see Supplemental Information). This differential gene expression signature was used in Ingenuity Pathway Analysis (IPA) tool [15] as well as the open-source software ToppGene suite [12], both of which revealed cell movement and adhesion signatures as top Gene Ontology process. Furthermore, IPA analysis revealed actin cytoskeleton and RhoA signaling as critical pathways.

For deriving the cell movement, Actin cytoskeleton, and RhoA signaling signatures, we extracted the differentially expressed genes in these processes and pathways from IPA and used the raw RNA-seq counts to get the average expression. The values were RMS normalized across LH-CIL and H-CIL values and log_2_ transformed and plotted using the MeV software [11].

## Supporting information

Correction to the original manuscript

Other files - not used in the main text

Figures in the manuscript

Codes and data

## Acknowledgments

We acknowledge Applied Bioinformatics Labs and Genome Technology Center at New York University Langone Health for providing the necessary resources for this project. Also, we would like to acknowledge Dr. Adriana Heguy, Division of Advanced Research Technologies at New York University Langone Health, Dr. Aris Tsirigos, Applied Bioinformatics Labs and Dr. David Fenyo, Department of Biochemistry and Molecular Pharmacology for their support.

## Supplemental Information

query.png: Contains the original query used in the GDC portal to download the images

Gibbs_Models.pdf: Contains a brief description of the Gibbs processes

### Supplemental Data

Supplemental Data 01: Supplemental_Data_01_Geyer_Stats, Geyer model statistics

Supplemental Data 02: Supplemental_Data_02_Clusters, Cluster 1 and Gibbs cluster association for the patients

Supplemental Data 03: Supplemental_Data_03_Indices, Optimal number of clusters indicies by NbClust

Supplemental Data 04: Supplemental_Data_04_Centroids, Convergence of the Gibbs process - centroids information

Supplemental Data 05: Supplemental_Data_05_Survival, Survival information

Supplemental Data 06: Supplemental_Data_06_H-CIL_LH-CIL, information on LH-CIL and H-CIL division

Supplemental Data 07: Supplemental_Data_07_H-CIL-RNA, Original RNA-seq counts for H-CIL patients

Supplemental Data 08: Supplemental_Data_08_LH-CIL-RNA, Original RNA-seq counts for LH-CIL patients

Supplemental Data 09: Supplemental_Data_09_DGE, Differentially expressed genes between H-CIL and LH-CIL patients

Supplemental Data 10: Supplemental_Data_10_Cell_Movement, Implicated cell movement genes

Supplemental Data 11: Supplemental_Data_11_Cell_Movement_Average, Cell movement genes RNA-seq values (average and transformed)

Supplemental Data 12: Supplemental_Data_12_Actin_Pathway_Genes, Implicated Actin pathway genes

Supplemental Data 13: Supplemental_Data_13_Actin_Pathway_Average, Actin pathway RNA-seq values (average and transformed)

Supplemental Data 14: Supplemental_Data_14_RhoA_Pathway_Genes, Implicated RhoA pathway genes

Supplemental Data 15: Supplemental_Data_15_RhoA_Pathway_Average, RhoA pathway RNA-seq values (average and transformed)

### Supplemental Figures

Supplemental Figure 01: Shows time point/indices at which homotypic CIL is established in the Gibbs cluster

Supplemental Figure 02: Decrease in cell movement as revealed by IPA

Supplemental Figure 03: Shows 48/86 genes have differential expression consistent with the direction of decrease in cell movement

Supplemental Figure 04: Cell movement as the top GO process as revealed by Toppgene, an open source pathway analysis tool

Supplemental Figure 05: Altered RhoA signaling pathway

Supplemental Figure 06: Altered Actin Cytoskeleton pathway

Supplemental Figure 07: 86 cell movement gene signature heatmap

## Data availability

The downloaded TCGA images are yet to be submitted to a repository that can hold large imaging data (they are also available from the GDC portal). Codes and data for this project can be found at:

https://github.com/kannan-kasthuri/kannan-kasthuri.github.io/tree/master/Research/Gibbs

Figures with better resolution are supplied with the ‘Figures’ folder.

## References

[1] E. Rubak A. Baddeley and R. Turner. Spatial Point Patterns: Methodology and Applications with R. London: Chapman and Hall/CRC Press, 2015. ISBN: 9781482210200.

[2] Heaysman J.E.M. Abercrombie M. “Observations on the social behaviour of cells in tissue culture: I. Speed of movement of chick heart fibroblasts in relation to their mutual contacts.” In: Experimental Cell Research 5 (1953), pp. 111–131.

[3] Karthauser H.M. Abercrombie M. Heaysman J.E.M. “Social behaviour of cells in tissue culture. III. Mutual influence of sarcoma cells and fibroblasts”. In: Experimental Cell Research 13(2) (1957), pp. 276–291.

[4] Abercrombie. M. “Contact inhibition and malignancy”. In: Nature 281(5729) (1979), pp. 259–262.

[5] Adrian Baddeley and Rolf Turner. “Spatstat: an R package for analyzing spatial point patterns.” In: Journal of Statistical Software 12.6 (2005). ISSN 1548-7660, pp. 1–42. URL: www.jstatsoft.org.

[6] Brenner, David J., Shuryak, Igor and Sachs, Rainer K. Radiotherapy-Induced Carcinogenesis and Leukemogenesis: Mechanisms and Quantitative Modeling. Springer Berlin Heidelberg, 2014, pp. 205–226.

[7] Charlotte M. Jones-Todd, Peter Caie, Janine Illian, Ben C. Stevenson, Anne Savage, David J. Harrison, James L. Bown. “Unusual structures inherent in point pattern data predict colon cancer patient survival”. In: arXiv:1705.05938v1 [stat.ME] (2017).

[8] Malika Charrad et al. “NbClust: An R Package for Determining the Relevant Number of Clusters in a Data Set”. In: Journal of Statistical Software 61.6 (2014), pp. 1–36.

[9] Colaprico A, Silva TC, Olsen C, Garofano L, Cava C, Garolini D, Sabedot T, Malta TM, Pagnotta SM, Castiglioni I, Ceccarelli M, Bontempi G, Noushmehr H. “TCGAbiolinks: An R/Bioconductor package for integrative analysis of TCGA data.” In: Nucleic Acids Research (2015).

[10] Fife CM, McCarroll JA, Kavallaris M. “Movers and shakers: cell cytoskeleton in cancer metastasis”. In: Br. J. Pharmacol. 171(24) (2014), pp. 5507–5523.

[11] Eleanor Howe et al. “MeV: MultiExperiment viewer”. In: Mar. 2010, pp. 267–277.

[12] Jing Chen, Eric E. Bardes, Bruce J. Aronow, Anil G. Jegga. “ToppGene Suite for gene list enrichment analysis and candidate gene prioritization”. In: Nucleic Acids Research 37(2) (2009), W305–W311.

[13] Jing, H., Yang, Y., and Nishikawa, R. M. “Detection of clustered microcalcifications using spatial point process modeling.” In: Physics in Medicine and Biology 56(1) (2011), pp. 1–17.

[14] King, Benjamin A. et al. “Spatiotemporal Patterns of Tumor Occurrence in Children with Intraocular Retinoblastoma.” In: PLoS ONE 10(7) (2015), e0132932.

[15] Krämer A, Green J, Pollard J Jr, Tugendreich S. “Causal analysis approaches in Ingenuity Pathway Analysis.” In: Bioinformatics. 30(4) (2013), pp. 523–530.

[16] Leininger, T. J. “Bayesian analysis of spatial point patterns”. In: PhD dissertation (2014).

[17] Liu L, He B, Liu WM, Zhou D, Cox JV, Zhang XA. “Tetraspanin CD151 promotes cell migration by regulating integrin trafficking.” In: J Biol Chem. 282(43) (2007), pp. 31631–42.

[18] M. Abercrombie, E. J. Ambrose. “The Surface Properties of Cancer Cells: A Review”. In: Cancer Research 22(5) (1962), pp. 525–548.

[19] M. Abercrombie, J.E.M. Heaysman, H.M. Karthauser. “Social behaviour of cells in tissue culture. III. Mutual influence of sarcoma cells and fibroblasts.” In: Exp. Cell Res. 13(2) (1957), pp. 276–291.

[20] Mayor R, Carmona-Fontaine C. “Keeping in touch with contact inhibition of locomotion”. In: Trends Cell Biol. 20(6) (2010), pp. 319–328.

[21] McCarthy, J. D, Chen, Yunshun, Smyth, K. G. “Differential expression analysis of multifactor RNA-Seq experiments with respect to biological variation.” In: Nucleic Acids Research 40(10) (2012), pp. 4288–4297.

[22] Artem S. Novozhilov, Georgy P. Karev, and Eugene V. Koonin. “Biological applications of the theory of birth-and-death processes”. In: Briefings in Bioinformatics 7(1) (2006), pp. 70–85.

[23] Hisashi Ohtsuki and Hideki Innan. “Allele Frequency Spectrum in a Cancer Cell Population”. In: bioRxiv (2017).

[24] Paolo Vineis, Arthur Schatzkin and John D. Potter. “Models of carcinogenesis: an overview.” In: Carcinogenesis 31(10) (2010), pp. 1703–1709.

[25] Parish, Roger W., Schmidhauser, Christian, Schmidt, Thomas, Dudler, Robert K. “Mechanisms of tumour cell metastasis”. In: J Cell Sci Suppl. 8 (1987), pp. 181–197.

[26] Peter Bankhead, Maurice B. Loughrey, José A. Fernández, Yvonne Dombrowski, Darragh G. McArt, Philip D. Dunne, Stephen McQuaid, Ronan T. Gray, Liam J. Murray, Helen G. Coleman, Jacqueline A. James, Manuel Salto-Tellez and Peter W. Hamilton. “QuPath: Open source software for digital pathology image analysis.” In: Sci. Rep. 7(16878) (2017).

[27] C.J. Preston. “Spatial birth-and-death processes”. In: Bulletin of the International Statistical Institute 46 (1977), pp. 371–391.

[28] B. D. Ripley. “Modelling Spatial Patterns”. In: Journal of the Royal Statistical Society. Series B 39(2) (1977), pp. 172–212.

[29] Mayor R. Roycroft A. “Molecular basis of contact inhibition of locomotion.” In: Cell Mol Life Sci. 73(6) (2015), pp. 1119–1130.

[30] Saribudak, A., Kucharavy, H., Hubbard, K., and Uyar, M. U. “Spatial Heterogeneity Analysis in Evaluation of Cell Viability and Apoptosis for Colorectal Cancer Cells.” In: IEEE Journal of Translational Engineering in Health and Medicine 4 (2016), p. 4300209.

[31] C.M. Thompson and L. Shure. Image Processing Toolbox: For Use with MATLAB;[user’s Guide]. MathWorks, 1995.

[32] Tuck AB, Hota C, Wilson SM, Chambers AF. “Osteopontin-induced migration of human mammary epithelial cells involves activation of EGF receptor and multiple signal transduction pathways.” In: Oncogene. 22(8) (2003), pp. 1198–205.

[33] van den Berg YW, van den Hengel LG, Myers HR, Ayachi O, Jordanova E, Ruf W, Spek CA, Reitsma PH, Bogdanov VY, Versteeg HH. “Alternatively spliced tissue factor induces angiogenesis through integrin ligation.” In: Proc Natl Acad Sci USA. 106(46) (2009), pp. 19497–502.

[34] Wang J, Guan E, Roderiquez G, Calvert V, Alvarez R, Norcross MA. “Role of tyrosine phosphorylation in ligand-independent sequestration of CXCR4 in human primary monocytes-macrophages.” In: J Biol Chem. 276(52) (2001), pp. 49236–43.

[35] Ann P. Wheeler and Anne J. Ridley. “Why three Rho proteins? RhoA, RhoB, RhoC, and cell motility”. In: Experimental Cell Research 301.1 (2004), pp. 43–49.

[36] M. Zhou et al. “Radiomics in Brain Tumor: Image Assessment, Quantitative Feature Descriptors, and Machine-Learning Approaches”. In: American Journal of Neuroradiology (2017). ISSN: 0195-6108.

[37] Zimmermann, Juliane and Camley, Brian A. and Rappel, Wouter-Jan and Levine, Herbert. “Contact inhibition of locomotion determines cell-cell and cell-substrate forces in tissues”. In: Proceedings of the National Academy of Sciences 113(10) (2016), pp. 2660–2665.

