## Supplementary material for "Gibbs Process Determines Survival and Reveals Contact-Inhibition Genes in Glioblastoma Multiforme": Correction to the original manuscript

In page $3$, the line "That is, if the cells are contact-inhibited, we can expect the fitted Gibbs hard-core constraint, $G_r$, to be approximately the same as $L_r$, indicating the model fits the statistics as determined by the $L$-function."

should be replaced by,

"That is, if the cells are contact-inhibited, we can expect the fitted Gibbs hard-core constraint, $G_r$, to be approximately the same as the optimal value of $\mid L_r - r \mid$, indicating the model fits the statistics as determined by the $L$-function. Therefore, we will use this metric for our results. That is, $L_r \equiv \ \mid L_r - r \mid$."

***Note***: This corrigendum does not affect the narrative or the claims made in the manuscript since this is the metric that is used for computations, as it can be seen in the "recursiveFitGeyer_AIC.R" code that was supplied, especially the line:

```
discrep <- function(r) {
  return(abs(as.function(Lest(p))(r) - r))
}

res <- optimise(discrep, interval = c(0, 10), maximum = TRUE)
```
