## Supplementary material for "Gibbs Process Determines Survival and Reveals Contact-Inhibition Genes in Glioblastoma Multiforme": Gibbs_Models.pdf

The Gibbs point processes belong to a large class of point processes where interaction between the points are considered for modeling. The interactions can be repulsive or attractive depending on the type of model we choose to represent the physical system while the null model is typically a Poisson point process [1].

**Strauss Process**

Strauss process is a generalization of the Gibbs process and can be considered as one of the variations of the basic Gibbs hard core process. In the Gibbs hard core process, it is physically impossible for two points to lie closer than $h$ distance apart. But if those points are not impossible but *improbable* the model is called the Strauss process. Therefore, in Strauss process, close pair of points are penalized instead of forbidden. Thus, if the probability density of the Gibbs hard core process is given by,

$$f(\mathbf{x}) = \begin{cases} \beta & \text{if hard core constraint is satisfied,} \\ 0 & \text{if hard core constraint is violated.} \end{cases}$$

where $\beta$ is the 'abundance' defined in the "Overview of the Gibbs process and L-function" section; then probability density for the Strauss process is,

$$f(\mathbf{x}) = \mathbf{c}\,\beta\,\gamma^{\mathbf{t(x,h)}}$$

where $c$ is a constant, the parameter $0 \leq \gamma \leq 1$ is the interaction parameter, and $t(\mathbf{x}, \mathbf{h})$ is the number of unordered cells which are closer than the interaction distance $h$.

Thus, according to above definition of probability density, a collection of cells with many pairs of close cells is much less likely than a collection of cells with only a few close pairs.

**Geyer Saturation Model**

We note that in the above Strauss process, if $\gamma = 1$, then it is essentially a Poisson point process and $\gamma > 1$ can introduce clustering. However, $\gamma > 1$ cannot exist because the probability density is not integrable since it can take arbitrarily large values. Hence, to include the clustering phenomena, Geyer [2] suggested a *saturation model* where the probability density is,

$$f(\mathbf{x}) = \mathbf{c}\,\beta\,\gamma^{\mathbf{min(s,t(x,h))}}$$

where $c, \beta, t(\mathbf{x}, h)$ are as described above and $s \geq 0$ is the saturation parameter. Note that $s = 0$ is a Poisson process and if $s > 0$, the interaction $\gamma$ may take any positive value. If $\gamma < 1$, we will have an inhibiting phenotype and $\gamma > 1$ will have a clustering phenotype. Also, $s = \infty$ is a Strauss process. Thus, Geyer saturation model is a generalization of the Gibbs hard core process.

**References**


1. David Dereudre, Introduction to the theory of Gibbs point processes, arXiv:1701.08105v2 [math.PR]

2. C.J. Geyer. Likelihood inference for spatial point processes. In O.E. Barndorff-Nielsen, W.S. Kendall, and M.N.M. van Lieshout, editors, *Stochastic Geometry: Likelihood and Computation*. Number 80 in Monographs on Statistics and Applied Probability, Chapter 3, pages 79-140. Chapman and Hall/CRC. Boca Raton, FL, 1999.
