## Supplementary figures and images for "Gibbs Process Determines Survival and Reveals Contact-Inhibition Genes in Glioblastoma Multiforme"

### Fig.1.png

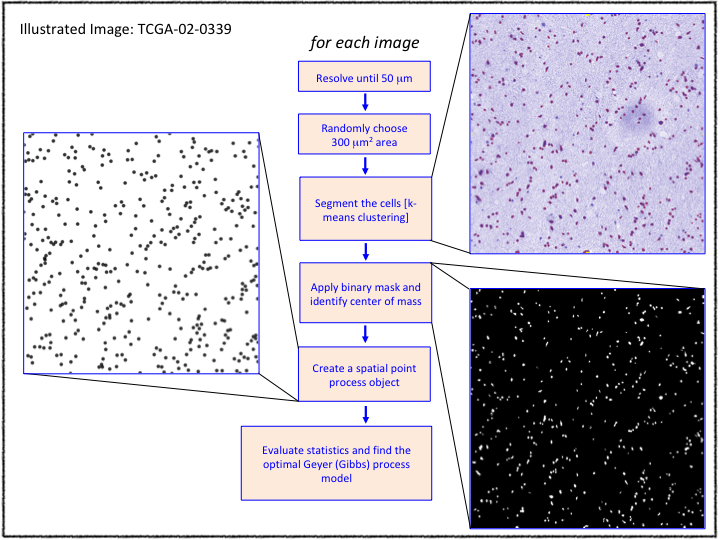

### Fig.2.png

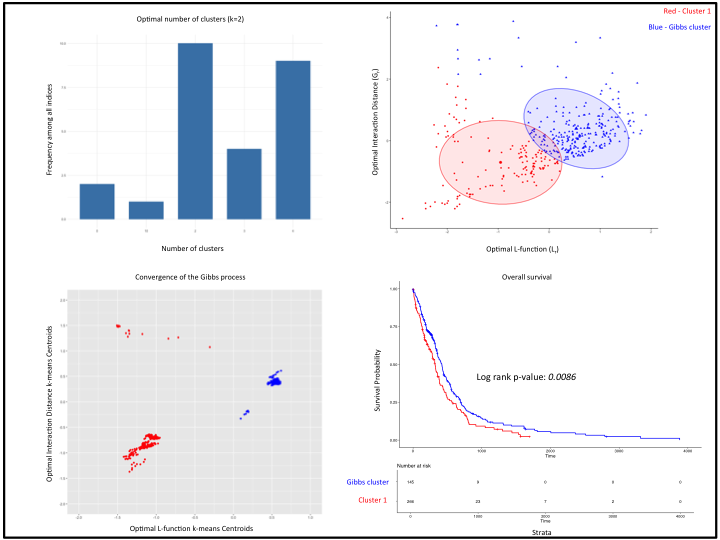

### Fig.3.png

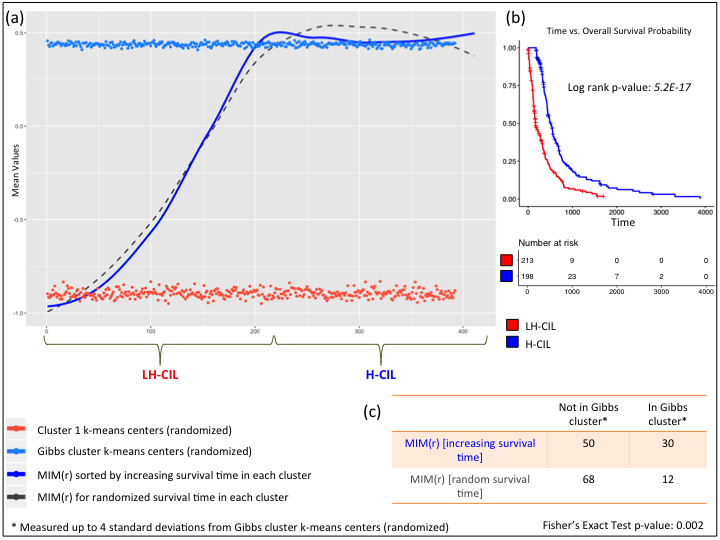

### Fig.4.png

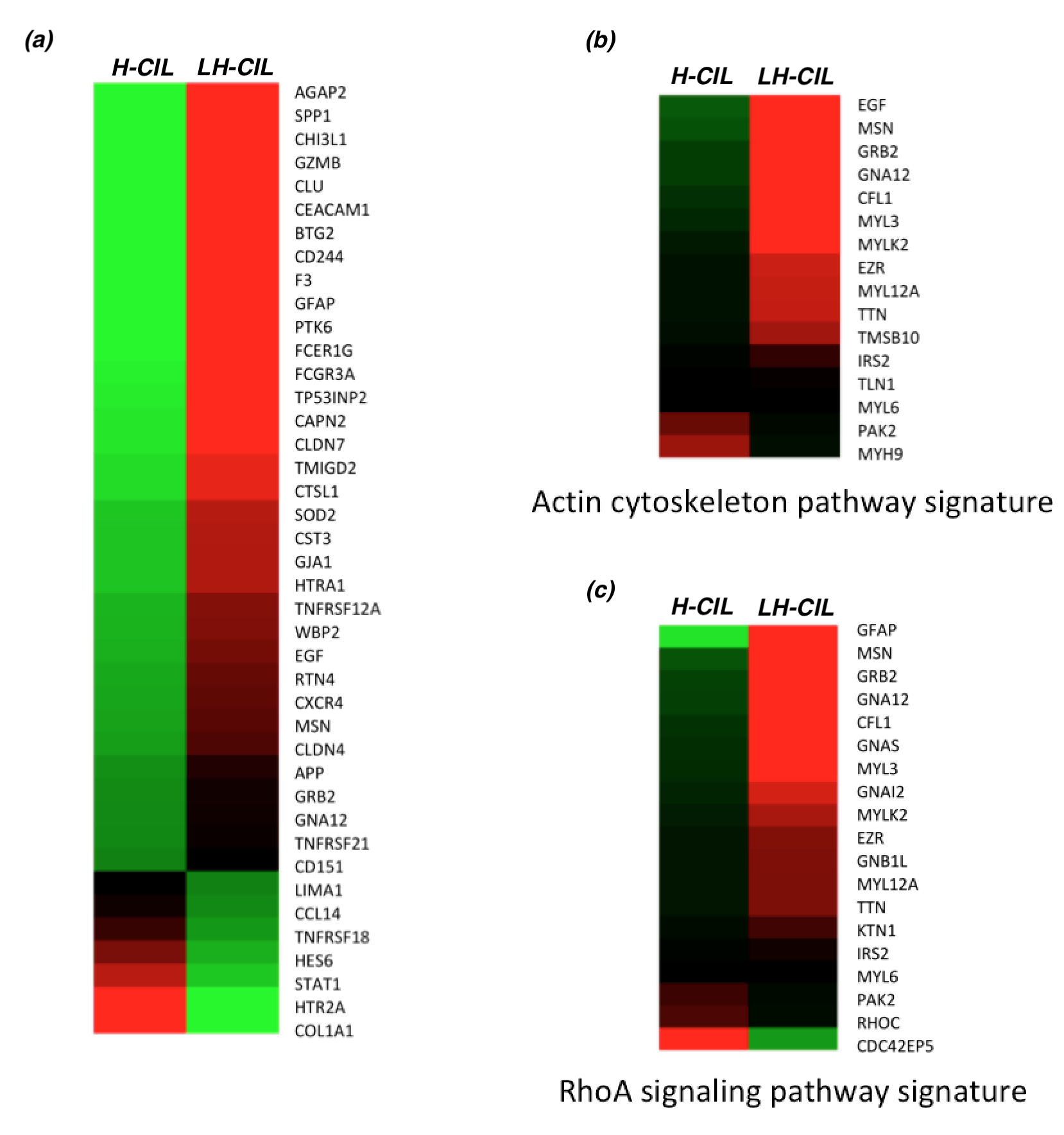

### query.png

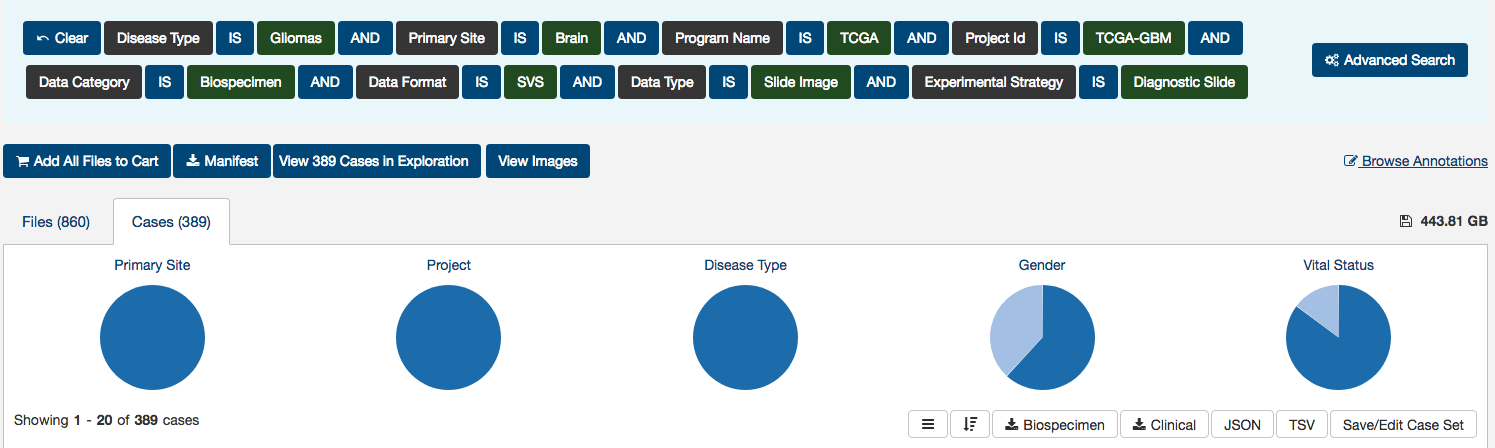

### Screen Shot 2019-04-12 at 12.19.50 PM.png

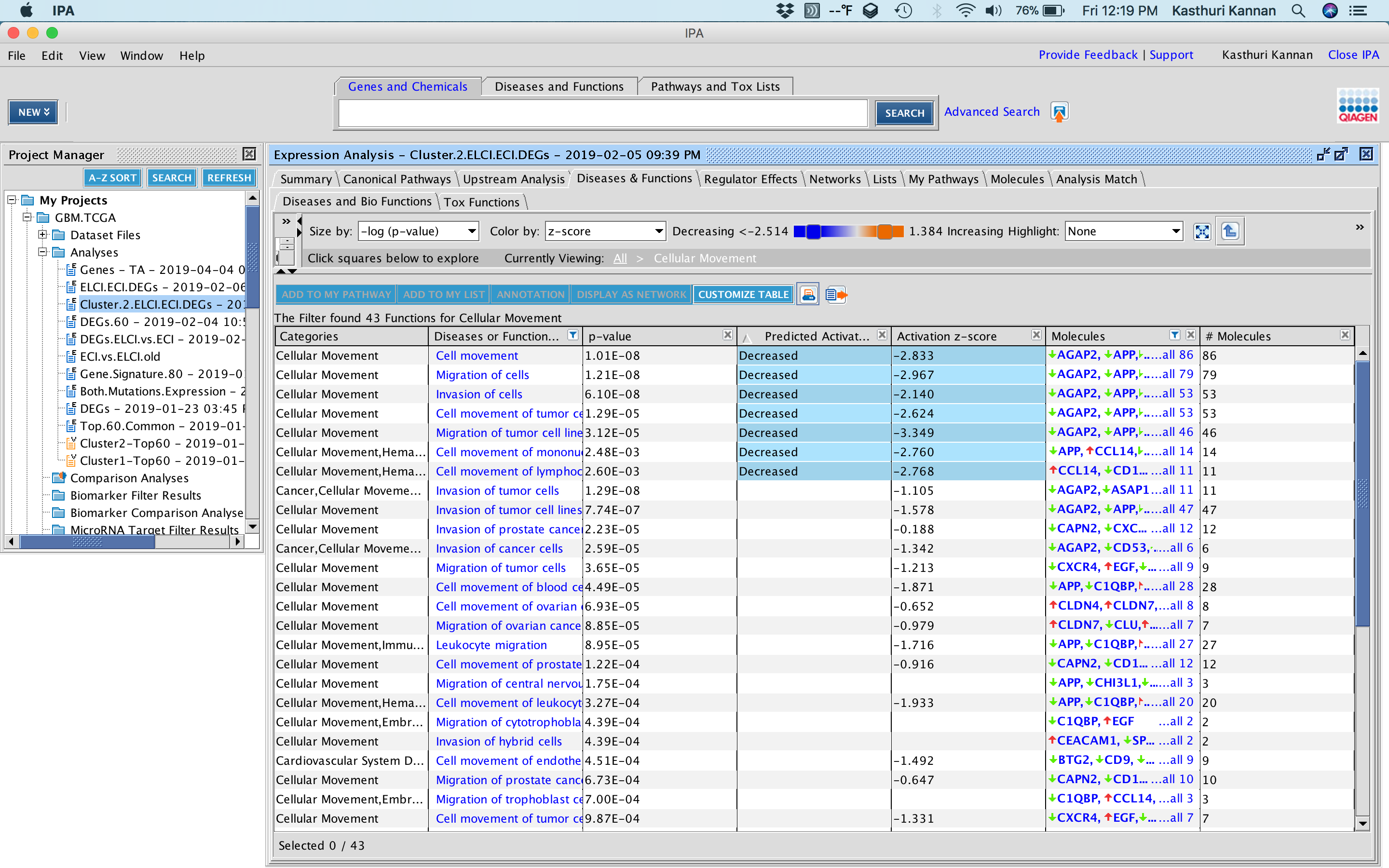

### Screen Shot 2019-04-12 at 12.20.12 PM.png

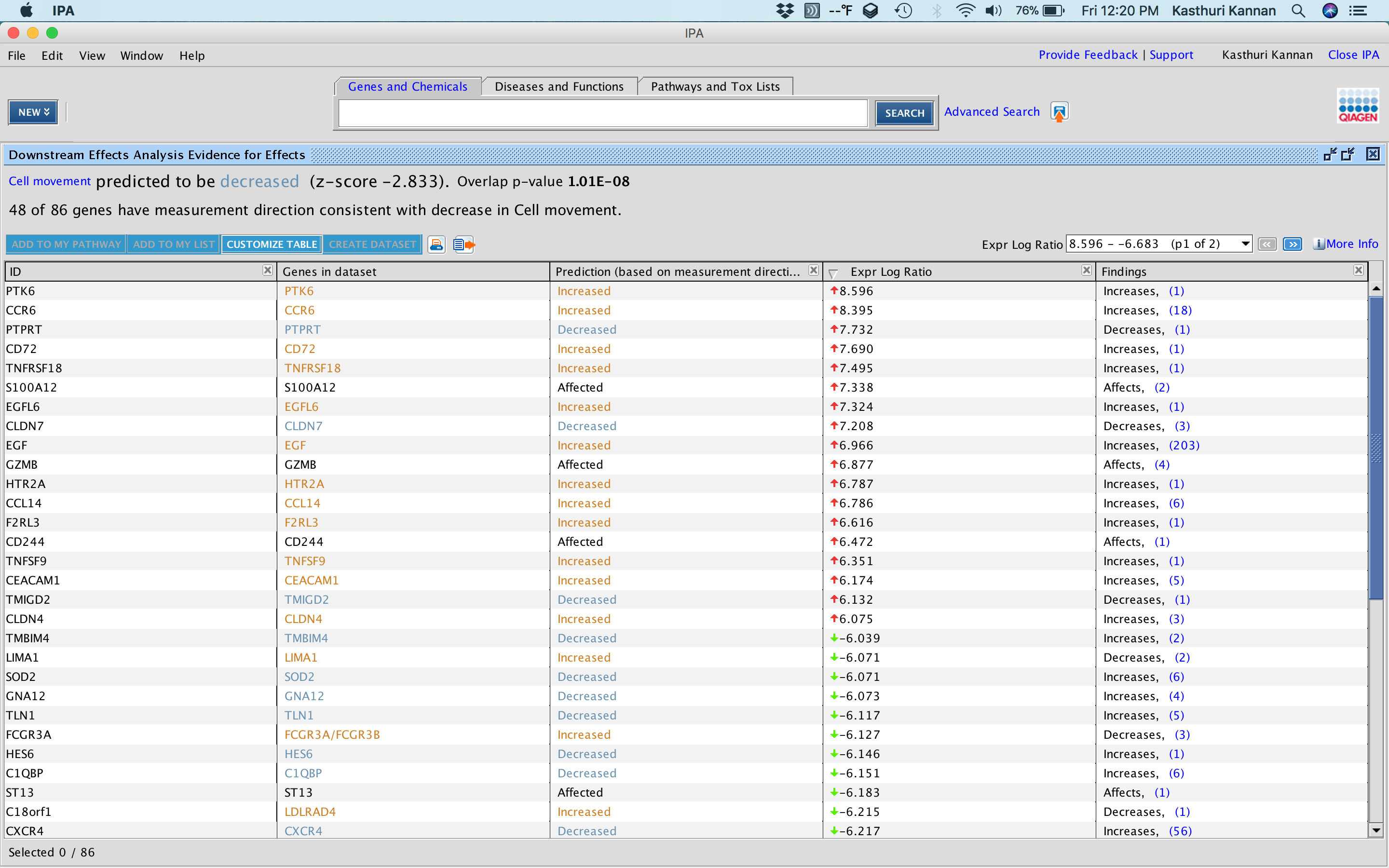

### Screen Shot 2019-04-12 at 12.28.01 PM.png

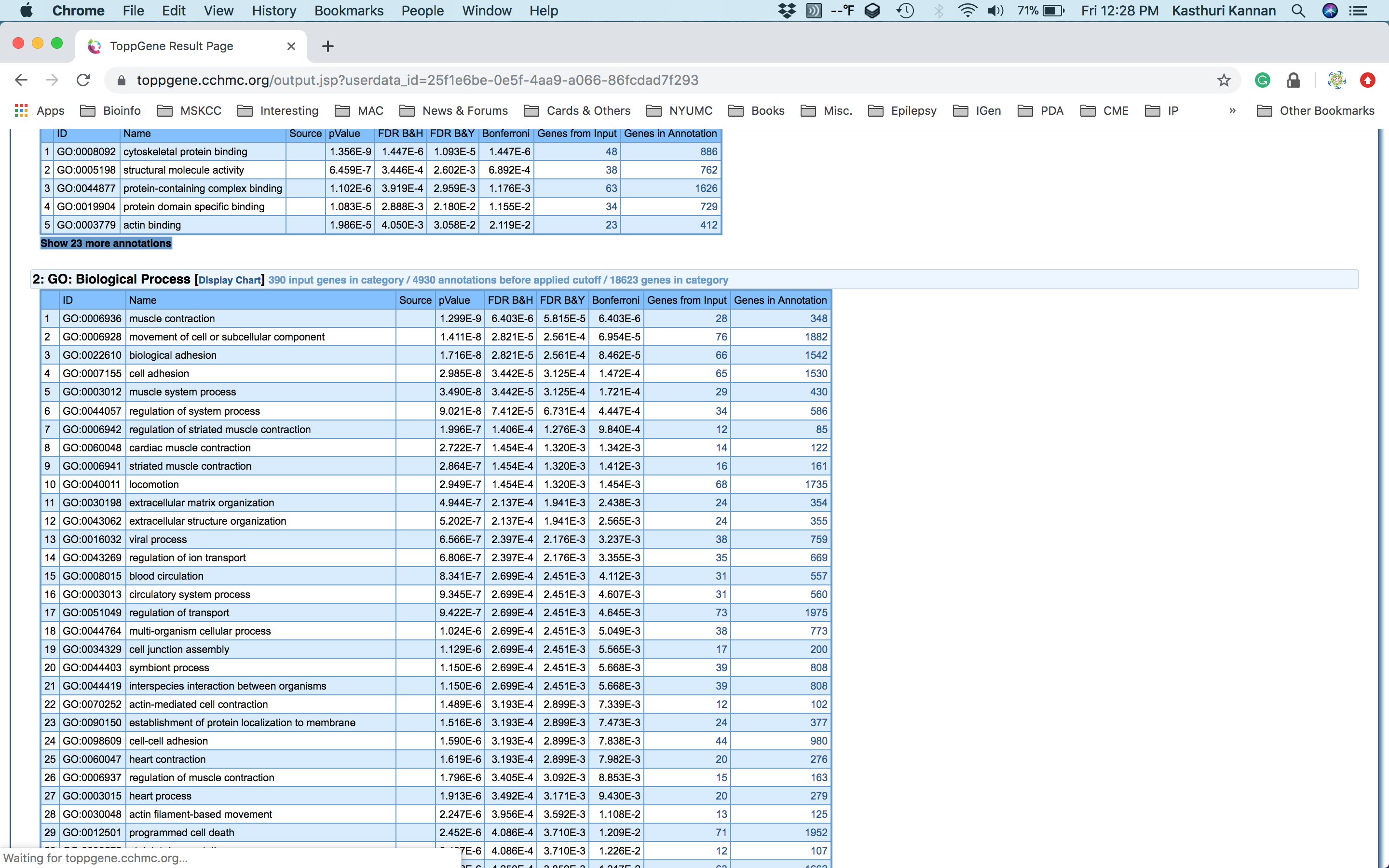

### Supplemental Figure 01.png

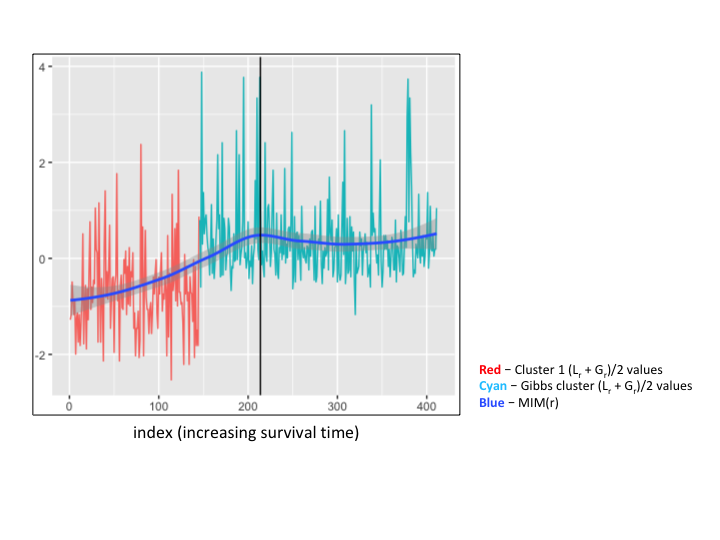

### Supplemental Figure 02.png

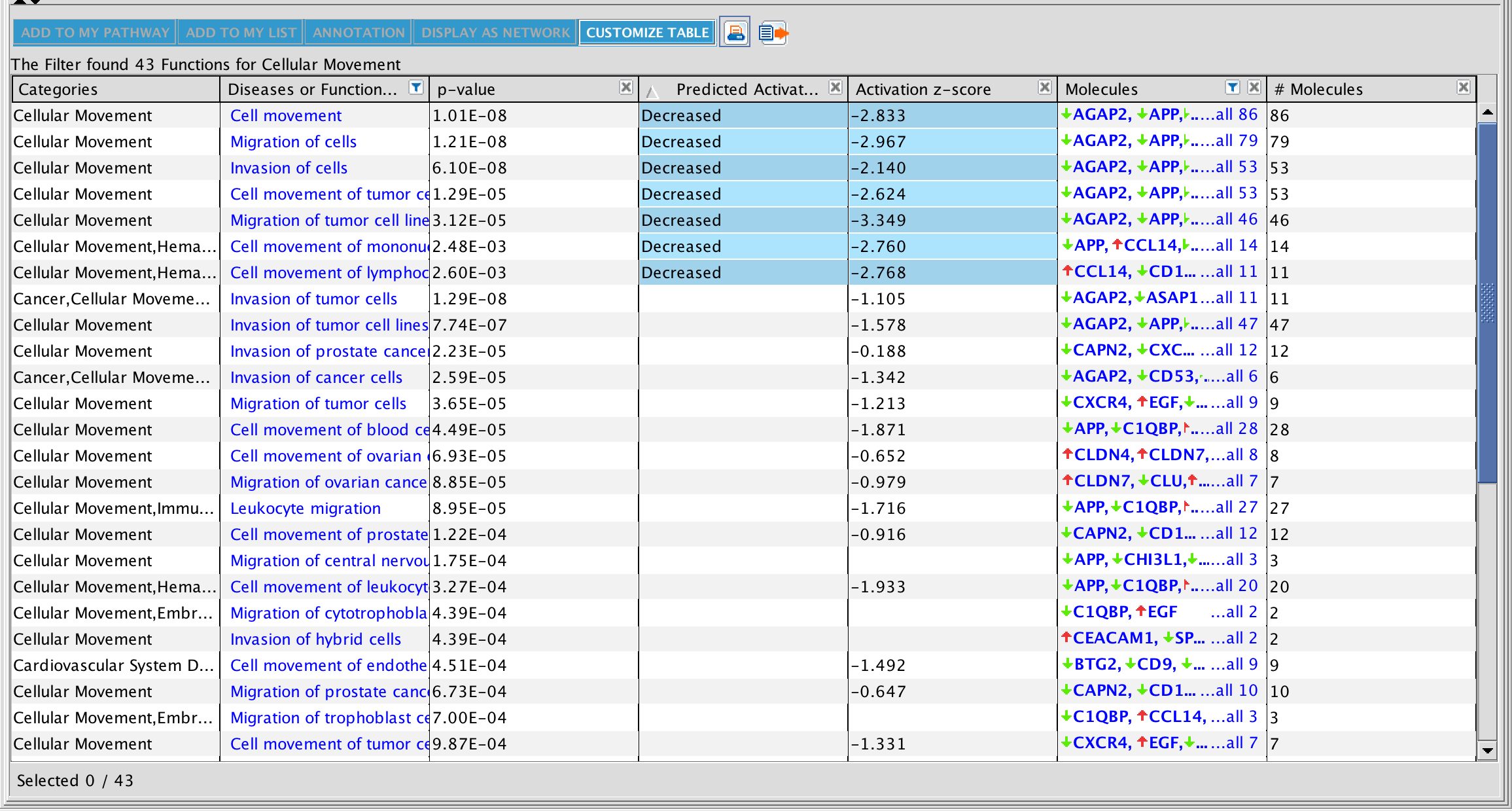

### Supplemental Figure 03.png

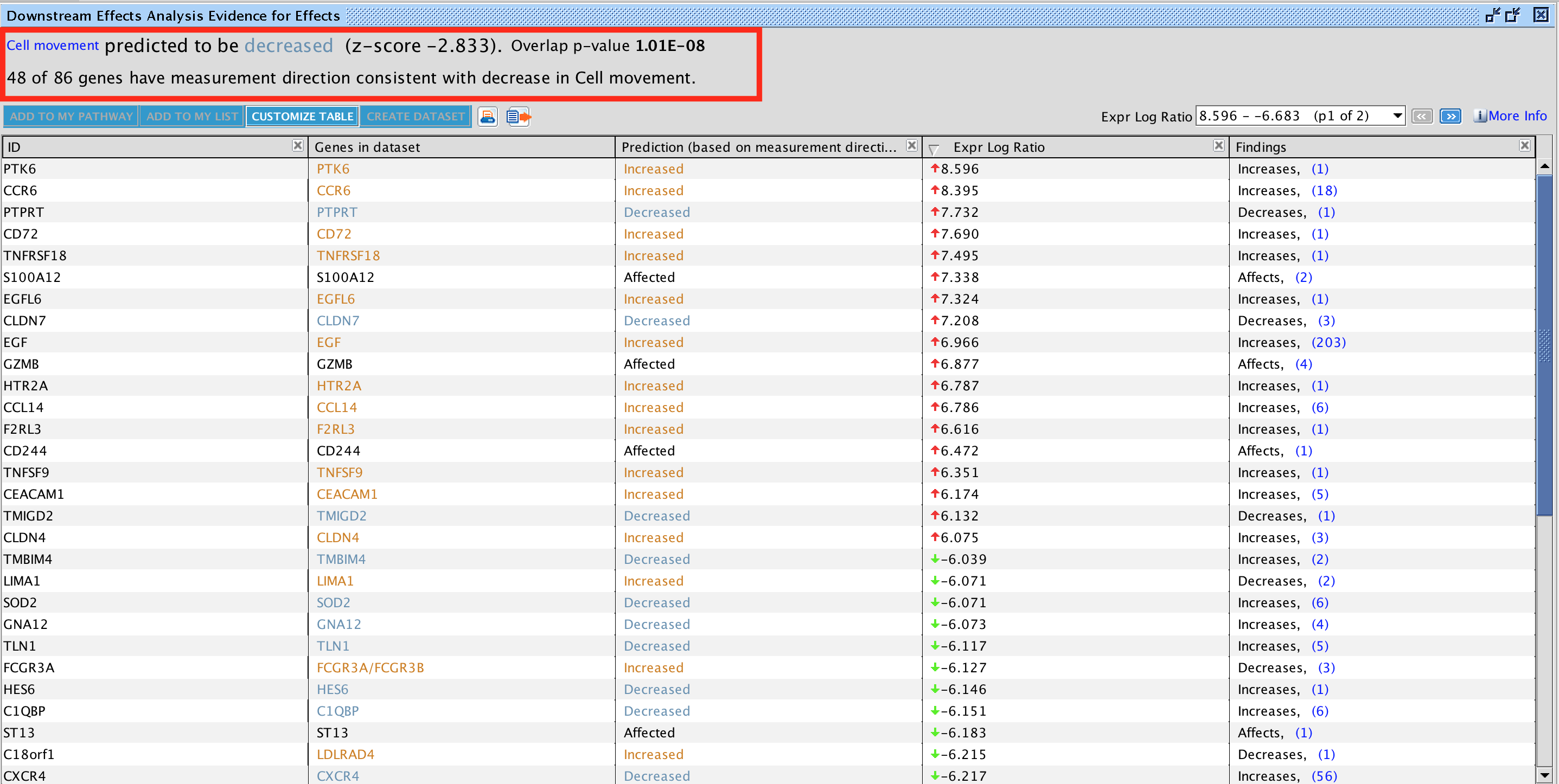

### Supplemental Figure 04.png

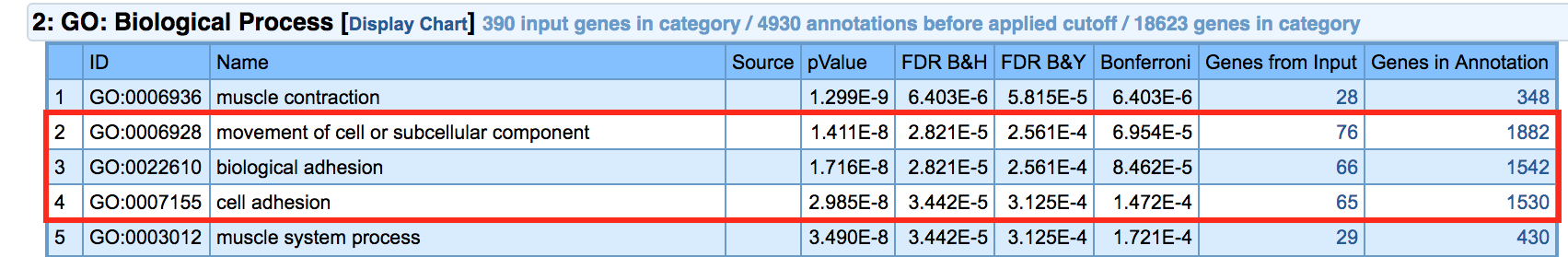

### Supplemental Figure 05.png

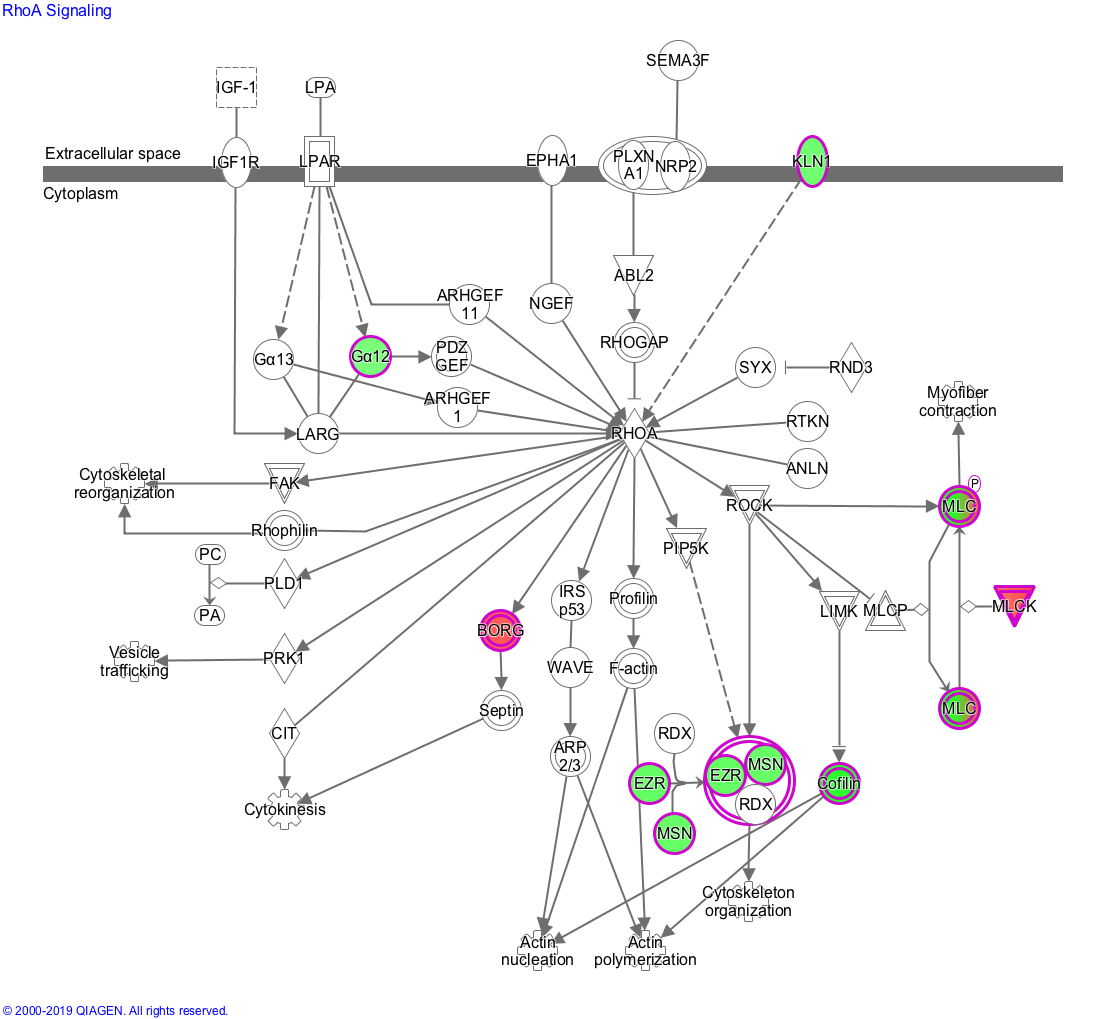

### Supplemental Figure 06.png

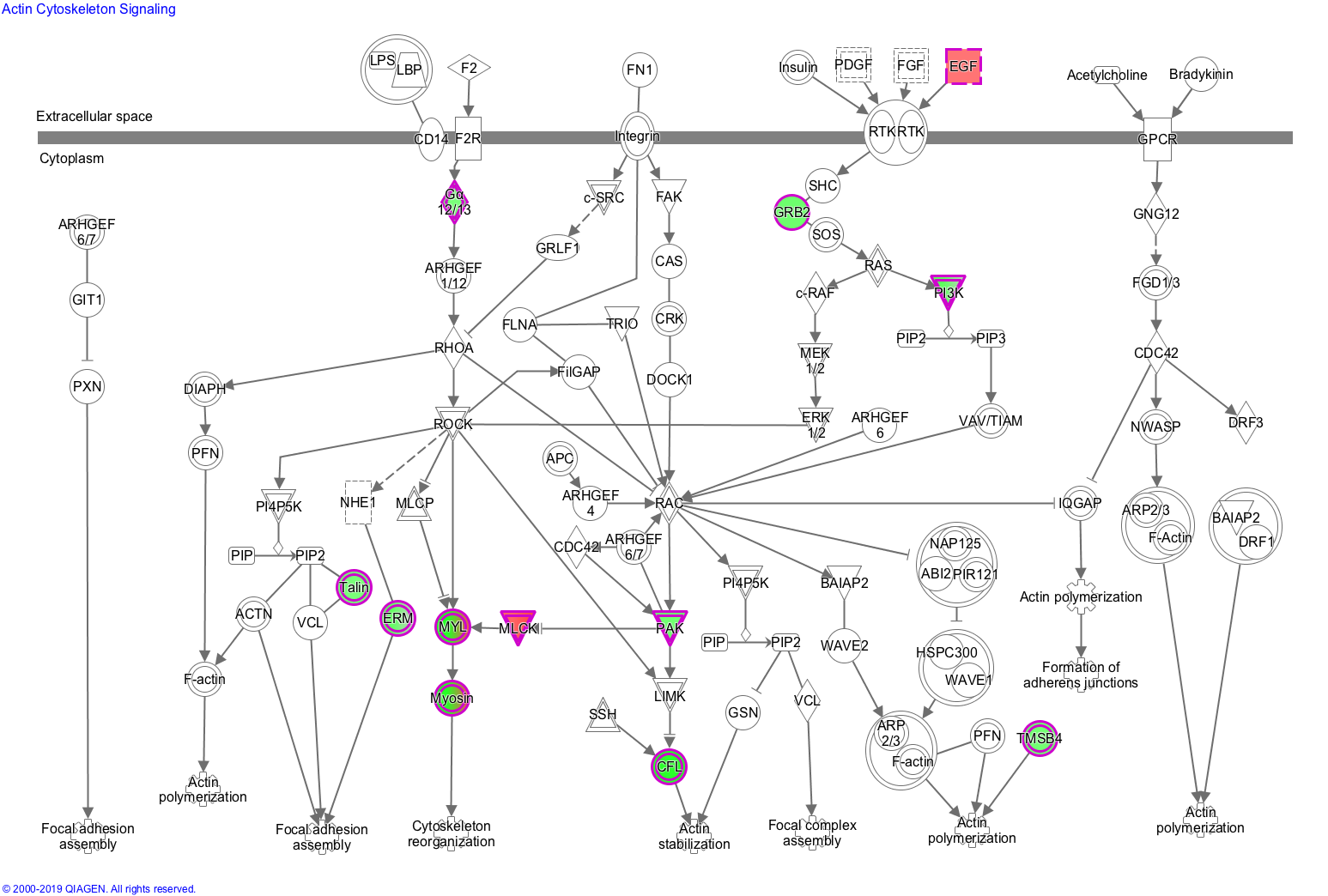

### Supplemental Figure 07.tiff

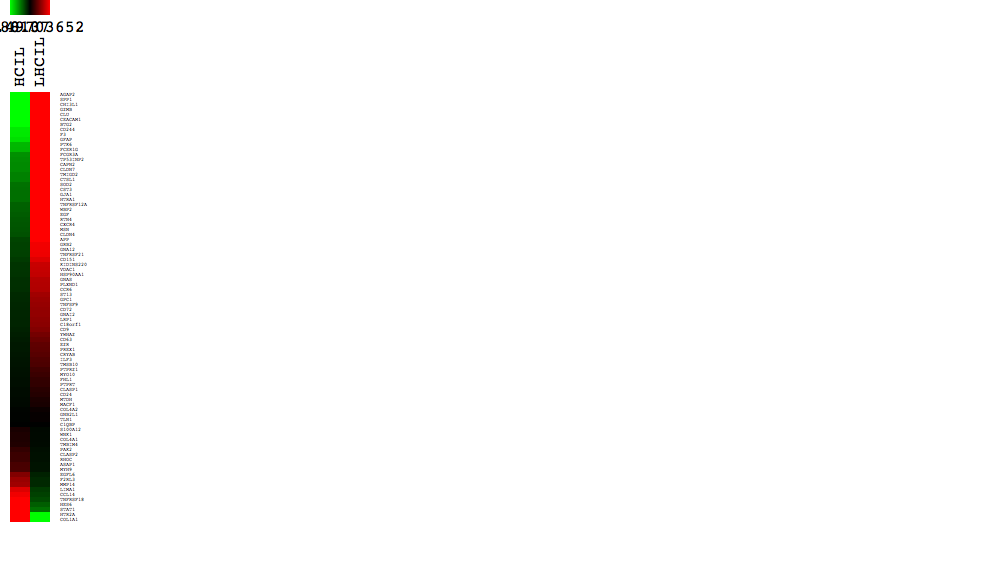
