## Supplementary material for "Gibbs Process Determines Survival and Reveals Contact-Inhibition Genes in Glioblastoma Multiforme": Other files - not used in the main text: Supplementary_Figure1.pptx

### Slide 1
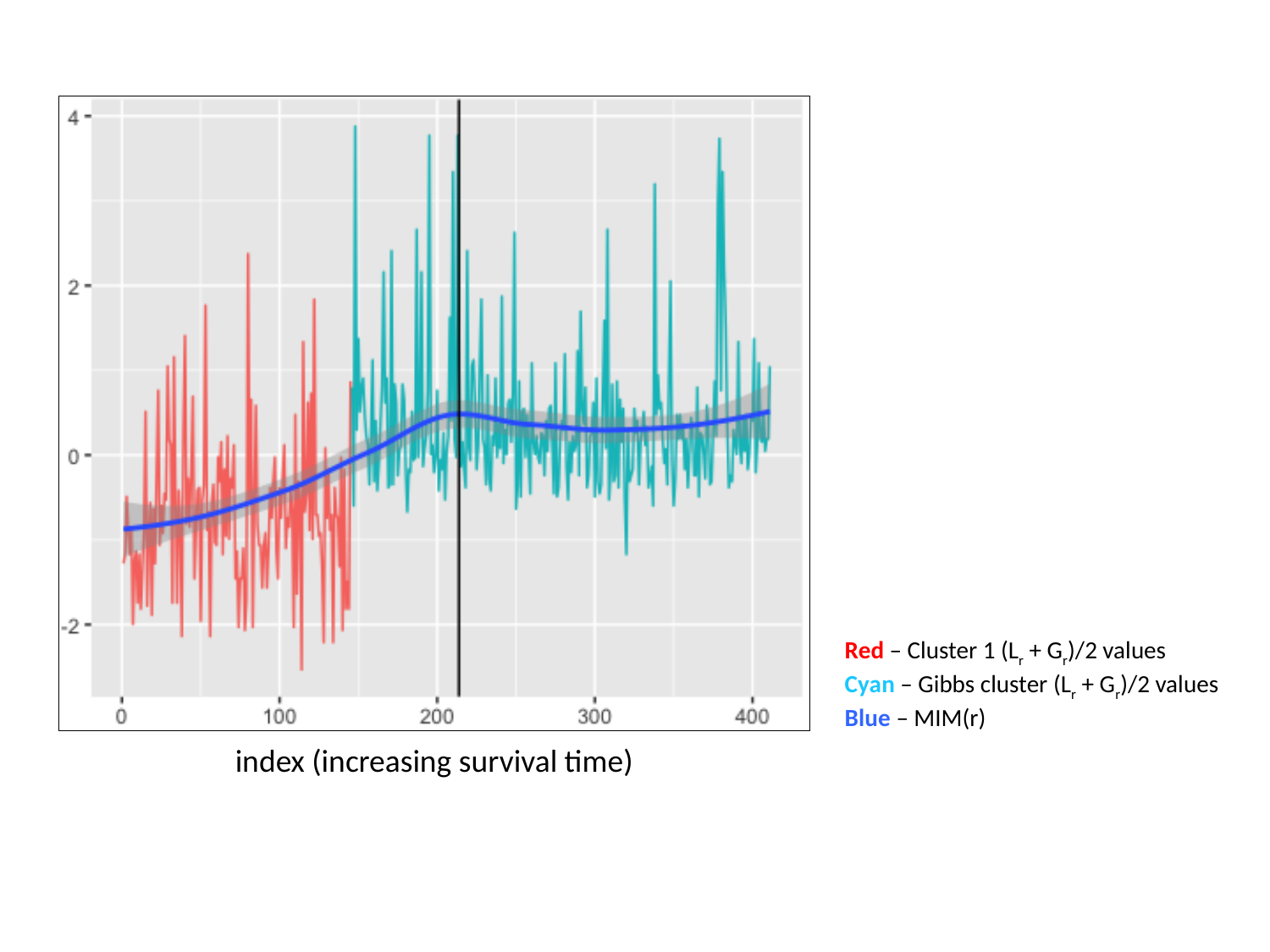

Red – Cluster 1 (L_r_ + G_r_)/2 values
Cyan – Gibbs cluster (L_r_ + G_r_)/2 values
Blue – MIM(r)
index (increasing survival time)
