## Supplementary material for "Gibbs Process Determines Survival and Reveals Contact-Inhibition Genes in Glioblastoma Multiforme": Codes and data: Figure.2.Code.R

```
# This code performs cluster analysis on the Geyer fitted model

# Load relevant libraries and set seed
library(cluster)
library(factoextra)
library(readxl)
library("NbClust")
library(survival)
library(survminer)
library(dplyr)
library("TCGAbiolinks")
library(RTCGA.clinical)

set.seed(123)


# Load the Geyer model data
geyer_data <- read_excel("~/Desktop/TCGA/Results-GBM/Classifier/GBM_Geyer.xlsx")

# Get only the relevent features for unsupervised clustering- gamma, optL, LObj, interaction distance
features <- geyer_data[,9:12]
features <- features %>% select(OptL,`Interaction Distance`)
features <- scale(features)
features <- na.omit(features)

# -------------------------------------------------------------------------------------------#
# Determining the optimal number of clusters and visualize
# -------------------------------------------------------------------------------------------#
res.nbclust <- NbClust(features, distance = "euclidean", min.nc = 2, max.nc = 10, method = "median", index ="all")
factoextra::fviz_nbclust(res.nbclust) + theme_minimal()

write.csv(res.nbclust$All.index, "~/Dropbox (NYU Langone Health)/my.research/spatial.analysis/Gibbs/Figure.02/optimal.k.index.csv")
write.csv(res.nbclust$All.CriticalValues, "~/Dropbox (NYU Langone Health)/my.research/spatial.analysis/Gibbs/Figure.02/optimal.k.all.critical.values.csv")
write.csv(res.nbclust$Best.nc, "~/Dropbox (NYU Langone Health)/my.research/spatial.analysis/Gibbs/Figure.02/optimal.k.best.nc.csv")
write.csv(res.nbclust$Best.partition, "~/Dropbox (NYU Langone Health)/my.research/spatial.analysis/Gibbs/Figure.02/optimal.k.best.partition.csv")

saveRDS(res.nbclust, file = "~/Dropbox (NYU Langone Health)/my.research/spatial.analysis/Gibbs/Figure.02/Optimal-k/res.nbclust.rds")

# -------------------------------------------------------------------------------------------#
# We now run k-means with the optimal number of clusters and visualize
# -------------------------------------------------------------------------------------------#
km.res <- kmeans(features, 2)
fviz_cluster(list(data = features, cluster = km.res$cluster), ellipse.type = "norm", geom = "point", stand = FALSE, ggtheme = theme_classic(),
             ellipse.level = 0.70, ellipse.alpha = 0.1, palette = c("red","blue")) +
  xlab("Optimal L-function") + ylab("Optimal Interaction Distance") 

write.csv(km.res$cluster,  "~/Dropbox (NYU Langone Health)/my.research/spatial.analysis/Gibbs/Figure.02/k-means/km.cluster.csv")
write.csv(km.res$centers,  "~/Dropbox (NYU Langone Health)/my.research/spatial.analysis/Gibbs/Figure.02/k-means/km.centers.csv")
write.csv(km.res$totss,  "~/Dropbox (NYU Langone Health)/my.research/spatial.analysis/Gibbs/Figure.02/k-means/km.totss.csv")
write.csv(km.res$withinss,  "~/Dropbox (NYU Langone Health)/my.research/spatial.analysis/Gibbs/Figure.02/k-means/km.withinss.csv")
write.csv(km.res$tot.withinss,  "~/Dropbox (NYU Langone Health)/my.research/spatial.analysis/Gibbs/Figure.02/k-means/km.tot.wininss.csv")
write.csv(km.res$betweenss,  "~/Dropbox (NYU Langone Health)/my.research/spatial.analysis/Gibbs/Figure.02/k-means/km.betweenss.csv")
write.csv(km.res$size,  "~/Dropbox (NYU Langone Health)/my.research/spatial.analysis/Gibbs/Figure.02/k-means/km.size.csv")
write.csv(km.res$iter,  "~/Dropbox (NYU Langone Health)/my.research/spatial.analysis/Gibbs/Figure.02/k-means/km.iter.csv")
write.csv(km.res$ifault,  "~/Dropbox (NYU Langone Health)/my.research/spatial.analysis/Gibbs/Figure.02/k-means/km.ifault.csv")

saveRDS(km.res, file = "~/Dropbox (NYU Langone Health)/my.research/spatial.analysis/Gibbs/Figure.02/k-means//km.res.rds")

# -------------------------------------------------------------------------------------------#
# We now plot cluster centers as we iterate through the samples to show convergence
# -------------------------------------------------------------------------------------------#

Get.Centroids <- function(i,cluster.centers, features)  {
  
  # Load the Geyer model data
  g <- read_excel("~/Desktop/TCGA/Results-GBM/Classifier/GBM_Geyer.xlsx")
  
  print(i)
  # Get the first 'i' patients
  g <- g[1:i,]
  
  # Get only the relevent features for unsupervised clustering- gamma, optL, LObj, interaction distance
  
  # features <- scale(geyer_data[,9:12])
  f <- g[,9:12]
  f <- f %>% select(OptL,`Interaction Distance`)
  f <- scale(f)
  f <- na.omit(f)

  # Run k-means for k = 2 and get the centroids
  res <- kmeans(na.omit(f), 2)
  centers.df <- res$centers[order(res$centers[,1]),]
  
  # We want to make them as a row vector with cluster 1 centroids followed by cluster 2 centroids
  centers.cluster.1 <- as.data.frame(t(centers.df[1,1:2]))
  centers.cluster.2 <- as.data.frame(t(centers.df[2,1:2]))
  
  # Append cluster 1 and cluster 2 centers
  centers.df <- cbind(centers.cluster.1,centers.cluster.2)
  cluster.centers <- rbind(cluster.centers, centers.df)
  
  return(cluster.centers)

}


cluster.centers <- data.frame(matrix(ncol = 4, nrow = 0))

for (i in seq(20, 411, by=1)) {
  
  print(paste0("Processing samples 1:", i))

  cluster.centers <- Get.Centroids(i,cluster.centers,f)
  
}

colnames(cluster.centers)[1] <- c("Cluster.1.OptL")
colnames(cluster.centers)[2] <- c("Cluster.1.IntD")
colnames(cluster.centers)[3] <- c("Cluster.2.OptL")
colnames(cluster.centers)[4] <- c("Cluster.2.IntD")

ggplot(data = cluster.centers, aes(x = Cluster.1.OptL, y = Cluster.1.IntD, color = "Cluster 1")) + geom_point() + 
  geom_point(mapping = aes(x = Cluster.2.OptL, y = Cluster.2.IntD, color = "Cluster 2")) + scale_color_manual(values = c('Cluster 1' = 'red', 'Cluster 2' = 'blue')) +
  xlab("Optimal L-function k-means centroid (L)") + ylab("Optimal Interaction Distance k-means centroid (R)") + 
  scale_x_continuous(breaks = c(-2,-1.5,-1,-0.5,0,0.5,1), limits = c(-2,1)) +
  scale_y_continuous(breaks = c(-2,-1.5,-1,-0.5,0,0.5,1,1.5,2), limits = c(-2,2))

write.csv(cluster.centers,  "~/Dropbox (NYU Langone Health)/my.research/spatial.analysis/Gibbs/Figure.02/centroids/cluster.centers.csv")
saveRDS(cluster.centers, file = "~/Dropbox (NYU Langone Health)/my.research/spatial.analysis/Gibbs/Figure.02/centroids/cluster.centers.rds")


# -------------------------------------------------------------------------------------------#
# Survival analysis
# -------------------------------------------------------------------------------------------#

# Read survival data from TCGA

GBM.survInfo <- survivalTCGA(GBM.clinical, extract.cols = c("admin.disease_code", 
                                                            "patient.vital_status", 
                                                            "patient.days_to_last_followup", 
                                                            "patient.days_to_death"))

survival.data <- read_excel("~/Dropbox (NYU Langone Health)/my.research/spatial.analysis/Gibbs/Figure.02/survival.analysis/survival.data.xlsx")

# Merge these two data sets
merged.data <- merge(survival.data, GBM.survInfo, by = "bcr_patient_barcode")

# Fit survival and save
fit <- survfit(Surv(times, patient.vital_status) ~ Cluster, data = merged.data)
survp <- ggsurvplot(fit, data = merged.data, risk.table = TRUE, pval=TRUE, palette = c("red", "blue"))
ggsave(file = "~/Dropbox (NYU Langone Health)/my.research/spatial.analysis/Gibbs/Figure.02/survival.analysis/survival.pdf", print(survp))
surv_pvalue(fit, data = merged.data)

write.csv(merged.data, "~/Dropbox (NYU Langone Health)/my.research/spatial.analysis/Gibbs/Figure.02/survival.analysis/survival.and.cluster.data.csv")
saveRDS(fit, file = "~/Dropbox (NYU Langone Health)/my.research/spatial.analysis/Gibbs/Figure.02/survival.analysis/fit.rds")

# -------------------------------------------------------------------------------------------#
```
