## Supplementary material for "Gibbs Process Determines Survival and Reveals Contact-Inhibition Genes in Glioblastoma Multiforme": Codes and data: Figure.3.Code.R

```
# This code is for generating figure 3
#----------------------------------------------------------------------------------------#
# Generate cluster centroids mean (L,R) data and smoothed plus smoothed randomized data
#----------------------------------------------------------------------------------------#

# Load relevant libraries and set seed
library(tidyverse)
set.seed(123)

# Read the increasing survival data and convert into factors
incr.sur.df <- read_excel("~/Desktop/TCGA/Results-GBM/gene.expression/Figure.03/data.xlsx")
incr.sur.df$idx <- as.numeric(row.names(incr.sur.df))
incr.sur.df$`Mean(Cluster)` <- factor(incr.sur.df$`Mean(Cluster)`)

# Find the means of InterD and OptL
# 'lr.mean' is the column we want for plotting
incr.sur.df$lr.mean <- rowMeans(cbind(incr.sur.df$`Mean(ScaledOptL)`, 
                                      incr.sur.df$`Mean(ScaledIDs)`))

#----------------------------------------------------------------------------------------#
# Randomized analysis
#----------------------------------------------------------------------------------------#

# Take each cluster in the increasing survival data
incr.sur.cluster.1 <- incr.sur.df %>% filter(`Mean(Cluster)` == '1')
incr.sur.cluster.2 <- incr.sur.df %>% filter(`Mean(Cluster)` == '2')

# Make 100 iterations of each cluster 'lr' mean column
iter <- 100

cluster.1.random.lr.means <- as.data.frame(replicate(iter, sample(incr.sur.cluster.1$lr.mean)))
cluster.2.random.lr.means <- as.data.frame(replicate(iter, sample(incr.sur.cluster.2$lr.mean)))

# And combine the cluster matrices (by appending rows) and make a matrix of randomized means
random.lr.iter.means <- rbind(cluster.1.random.lr.means, cluster.2.random.lr.means)
random.lr.iter.means$idx <- as.numeric(row.names(random.lr.iter.means))

# Smooth each column and get the smoothed data
smoothed.random.lr.iter.means <- data.frame(matrix(ncol = 0, nrow = 80))
smoothed.random.lr.iter.idx <- data.frame(matrix(ncol = 0, nrow = 80))

for (i in seq(1,iter,by=1)) {
  lr.data <- random.lr.iter.means[,i]
  p <- ggplot(data = random.lr.iter.means, aes(x = idx, y = lr.data)) + stat_smooth()
  gb <- ggplot_build(p)
  smoothed.random.lr.iter.means <- cbind(smoothed.random.lr.iter.means,gb$data[[1]]$y)
  smoothed.random.lr.iter.idx <- cbind(smoothed.random.lr.iter.idx,gb$data[[1]]$x)
}

# Find the means of the columns
random.lr.means <- as.data.frame(rowMeans(smoothed.random.lr.iter.means))
random.lr.idx <- as.data.frame(rowMeans(smoothed.random.lr.iter.idx))

# random.lr is the final data we want for plotting
random.lr <- cbind(random.lr.idx,random.lr.means)
colnames(random.lr)[1:2] <- c("idx","LRmeans")

#----------------------------------------------------------------------------------------#
# Cluster center analysis and plots
#----------------------------------------------------------------------------------------#

# Read the cluster centers
cluster.centers <- readRDS(file = "~/Desktop/TCGA/Results-GBM/gene.expression/Figure.03/cluster.centers.rds")
cluster.centers$idx <- as.numeric(row.names(cluster.centers))

# Find the (L,R) mean for the cluster centers
cluster.centers$cluster.1.lr.mean <- rowMeans(cluster.centers[,1:2])
cluster.centers$cluster.2.lr.mean <- rowMeans(cluster.centers[,3:4])

# Randomize the means to get the trend for cluster 1
cluster.centers.1.lr.mean <- cluster.centers$cluster.1.lr.mean
cluster.1.permuted <- as.data.frame(replicate(iter, sample(cluster.centers.1.lr.mean)))
cluster.1.permuted.lr.mean <- rowMeans(cluster.1.permuted)
# 'randomized.cluster.1.lr.mean' is the data we want for plotting
cluster.centers$randomized.cluster.1.lr.mean <- cluster.1.permuted.lr.mean

# Randomize the means to get the trend for cluster 2
cluster.centers.2.lr.mean <- cluster.centers$cluster.2.lr.mean
cluster.2.permuted <- as.data.frame(replicate(iter, sample(cluster.centers.2.lr.mean)))
cluster.2.random.lr.mean <- rowMeans(cluster.2.permuted)
# 'randomized.cluster.2.lr.mean' is the data we want for plotting
cluster.centers$randomized.cluster.2.lr.mean <- cluster.2.random.lr.mean

#----------------------------------------------------------------------------------------#
# Generate plot
#----------------------------------------------------------------------------------------#

ggplot(data = random.lr, se = FALSE) +
  geom_smooth(aes(x = idx, y = LRmeans, color = "Rand. survival"), se = FALSE, size = 1, linetype = "dashed") +
  geom_smooth(data = incr.sur.df, aes(x = idx, y = lr.mean, color = "Incr. suvival"), se = FALSE, size = 1.5) +
  geom_point(data = cluster.centers, aes(x = idx, y = randomized.cluster.2.lr.mean, color = "Gibbs cluster")) +
  geom_point(data = cluster.centers, aes(x = idx, y = randomized.cluster.1.lr.mean, color = "Cluster 1")) +
  scale_color_manual(values = c('Cluster 1' = 'tomato', 'Gibbs cluster' = 'dodgerblue', 'Incr. suvival' = 'blue',
                                'Rand. survival' = 'gray30')) + 
  xlab("indices") + ylab("mean (L,R) values") # +
  # theme(panel.grid.major = element_blank(), panel.grid.minor = element_blank())

#----------------------------------------------------------------------------------------#
# Fisher's exact test analysis
#----------------------------------------------------------------------------------------#

# Find the upper and lower bounds for cluster 2 randomized LR mean data
# We use four standard deviations
upper.bound <- mean(cluster.centers$randomized.cluster.2.lr.mean) + 4*sd(cluster.centers$randomized.cluster.2.lr.mean)
lower.bound <- mean(cluster.centers$randomized.cluster.2.lr.mean) - 4*sd(cluster.centers$randomized.cluster.2.lr.mean)

# Get the smoothed values using ggplot_build for the increasing survival data, i.e., original
# data
p1 <- ggplot(data = incr.sur.df, aes(x = idx, y = lr.mean)) + stat_smooth()
gb.p1 <- ggplot_build(p1)
smoothed.incr.sur.df <- as.data.frame(gb.p1$data[[1]]$y)
names(smoothed.incr.sur.df)[1] <- c("LR")
# Check how many points lie above the lower bound and below the upper bound
count.s.surv.data <- smoothed.incr.sur.df %>% filter(LR > lower.bound & LR < upper.bound) %>% count()

# Get the smoothed values using ggplot_build for the randomized survival data
p2 <- ggplot(data = random.lr, aes(x = idx, y = LRmeans)) + stat_smooth()
gb.p2 <- ggplot_build(p2)
smoothed.rand.data <- as.data.frame(gb.p2$data[[1]]$y)
names(smoothed.rand.data) <- c("LR")

# Check how many points lie above the lower bound and below the upper bound
count.s.rand.data <- smoothed.rand.data %>% filter(LR > lower.bound & LR < upper.bound) %>% count()

# Form the contingency table
# mat(1,1) - increasing survival not belonging to cluster 2
# mat(1,2) - increasing survival belonging to cluster 2
# mat(2,1) - random survival not belonging to cluster 2
# mat(2,2) - random survival belonging to cluster 2
mat <- matrix(c(80-count.s.surv.data,80-count.s.rand.data,count.s.surv.data,count.s.rand.data), nrow = 2)
mat
m <- matrix(unlist(mat), 2)
# We want to see if there is an association of increasing survival time with cluster 2
fisher.test(m)

#----------------------------------------------------------------------------------------#
# Make box-plots for the ELCI and ECI clusters
#----------------------------------------------------------------------------------------#

ELCI.ECI.df <- incr.sur.df %>% filter(Cluster == "ELCI" | Cluster == "ECI")

ggplot(data = ELCI.ECI.df, aes(x = Cluster, y = `Mean(ScaledOptL)`)) + 
  geom_boxplot(aes(colour = Cluster)) + 
  xlab("Gibbs Subclusters") + ylab("Optimal L-function") +
  scale_color_manual(values=c("darkgreen", "purple")) 

# -------------------------------------------------------------------------------------------#
# Survival analysis - Cluster 1 + ELCI vs. ECI
# -------------------------------------------------------------------------------------------#

# Read survival data from TCGA

GBM.survInfo <- survivalTCGA(GBM.clinical, extract.cols = c("admin.disease_code", 
                                                            "patient.vital_status", 
                                                            "patient.days_to_last_followup", 
                                                            "patient.days_to_death"))

survival.data <- read_excel("~/Desktop/TCGA/Results-GBM/gene.expression/Figure.03/survival.data.xlsx")

# Merge these two data sets
merged.data <- merge(survival.data, GBM.survInfo, by = "bcr_patient_barcode")
# Fit survival and save
fit <- survfit(Surv(times, patient.vital_status) ~ Group, data = merged.data)
survp <- ggsurvplot(fit, data = merged.data, risk.table = TRUE, pval=FALSE, palette = c("red", "blue"))
ggsave(file = "~/Desktop/TCGA/Results-GBM/gene.expression/Figure.03/survival.pdf", print(survp))
surv_pvalue(fit, data = merged.data)

write.csv(merged.data, "~/Desktop/TCGA/Results-GBM/gene.expression/Figure.03/survival.and.cluster.data.csv")
saveRDS(fit, file = "~/Desktop/TCGA/Results-GBM/gene.expression/Figure.03/fit.rds")
```
