## Supplementary material for "Gibbs Process Determines Survival and Reveals Contact-Inhibition Genes in Glioblastoma Multiforme": Codes and data: Figure.4.Rcode.R

```
# Download TCGA Expression Data to do the analysis

library(TCGAbiolinks)
library(SummarizedExperiment)
library(dplyr)
library(readxl)

# Read the excel sheet
df <- read_excel("~/Desktop/TCGA/Results-GBM/gene.expression/ELCI.vs.ECI/Figure.04/ELCI.ECI.Cluster.2.Sample.Sheet.xlsx")

Cluster.1.df <- df %>% filter(df$Cluster == 'ELCI')
Cluster.2.df <- df %>% filter(df$Cluster == 'ECI')

# Cluster.1.df <- read_excel("~/Desktop/TCGA/Results-GBM/gene.expression/Cluster.1.Sample.Sheet.xlsx")
# Cluster.2.df <- read_excel("~/Desktop/TCGA/Results-GBM/gene.expression/Cluster.2.Sample.Sheet.xlsx")

# Downloading the data

listSamples <- Cluster.1.df$`Sample ID`
query <- GDCquery(project = "TCGA-GBM", data.category = "Gene expression", data.type = "Gene expression quantification",
                  experimental.strategy = "RNA-Seq", platform = "Illumina HiSeq", file.type = "results", barcode = listSamples, legacy = TRUE)
GDCdownload(query)
RnaseqSE <- GDCprepare(query, summarizedExperiment = FALSE)
# Matrix.C1 <- assay(RnaseqSE,"raw_count")
Matrix.C1 <- RnaseqSE[,seq(1,ncol(RnaseqSE),3)]

listSamples <- Cluster.2.df$`Sample ID`
query <- GDCquery(project = "TCGA-GBM", data.category = "Gene expression", data.type = "Gene expression quantification",
                  experimental.strategy = "RNA-Seq", platform = "Illumina HiSeq", file.type = "results", barcode = listSamples, legacy = TRUE)
GDCdownload(query)
RnaseqSE <- GDCprepare(query,summarizedExperiment = FALSE)
# Matrix.C2 <- assay(RnaseqSE,"raw_count")
Matrix.C2 <- RnaseqSE[,seq(1,ncol(RnaseqSE),3)]

write.csv(Matrix.C1,"~/Desktop/TCGA/Results-GBM/gene.expression/ELCI.vs.ECI/Figure.04/Original.Matrix.ELCI.csv")
write.csv(Matrix.C2,"~/Desktop/TCGA/Results-GBM/gene.expression/ELCI.vs.ECI/Figure.04/Original.Matrix.ECI.csv")

# Data analysis

# normalization of genes
Norm.Mat.C1 <- TCGAanalyze_Normalization(tabDF = Matrix.C1, geneInfo =  geneInfo)
Norm.Mat.C2 <- TCGAanalyze_Normalization(tabDF = Matrix.C2, geneInfo =  geneInfo)

# quantile filter of genes
Mat.C1.Filt <- TCGAanalyze_Filtering(tabDF = Norm.Mat.C1, method = "quantile", qnt.cut =  0.25)
Mat.C2.Filt <- TCGAanalyze_Filtering(tabDF = Norm.Mat.C2, method = "quantile", qnt.cut =  0.25)

d1 <- dim(Mat.C1.Filt)
d2 <- dim(Mat.C2.Filt)
min.dim <- min(d1[1],d2[1])

Mat.C1.Filt <- Mat.C1.Filt[1:min.dim, ]
Mat.C2.Filt <- Mat.C2.Filt[1:min.dim, ]

# Write the filtered matrices to a file
# write.csv(Mat.C1.Filt,"~/Desktop/TCGA/Results-GBM/gene.expression/ELCI.vs.ECI/ELCI.Filtered.Expression.csv")
# write.csv(Mat.C2.Filt,"~/Desktop/TCGA/Results-GBM/gene.expression/ELCI.vs.ECI/ECI.Filtered.Expression.csv")

# Diff.expr.analysis (DEA)
dataDEGs <- TCGAanalyze_DEA(mat1 = Mat.C1.Filt, mat2 = Mat.C2.Filt, Cond1type = "Cluster.1", Cond2type = "Cluster.2", fdr.cut = 0.001, logFC.cut = 6, method = "glmLRT")

# DEGs table with expression values in normal and tumor samples
dataDEGsFiltLevel <- TCGAanalyze_LevelTab(dataDEGs,"Cluster.1","Cluster.2",Mat.C1.Filt,Mat.C2.Filt)

write.csv(dataDEGsFiltLevel, "~/Desktop/TCGA/Results-GBM/gene.expression/ELCI.vs.ECI/ELCI.ECI.DEGs.csv")

# Enrichment Analysis EA
# Gene Ontology (GO) and Pathway enrichment by DEGs list
Genelist <- rownames(dataDEGsFiltLevel)

system.time(ansEA <- TCGAanalyze_EAcomplete(TFname="DEA genes Cluster.1 Vs Cluster.2",Genelist))

# Enrichment Analysis EA (TCGAVisualize)
# Gene Ontology (GO) and Pathway enrichment barPlot


dev.copy(pdf, "~/Desktop/deleteIt.png")

TCGAvisualize_EAbarplot(tf = rownames(ansEA$ResBP), 
                        GOBPTab = ansEA$ResBP,
                        GOCCTab = ansEA$ResCC,
                        GOMFTab = ansEA$ResMF,
                        PathTab = ansEA$ResPat,
                        nRGTab = Genelist, 
                        nBar = 10)

while (!(is.null(dev.list()["RStudioGD"]))){dev.off()}

my.Big.Matrix.Filtered <- cbind(Mat.C1.Filt, Mat.C2.Filt)
TCGAvisualize_PCA(my.Big.Matrix.Filtered,dataDEGsFiltLevel, ntopgenes = 200)
```
