## Supplementary material for "Gibbs Process Determines Survival and Reveals Contact-Inhibition Genes in Glioblastoma Multiforme": Codes and data: FitGeyer.R

```
#------------------------------------------------------------------#
# Load relevant libraries
#------------------------------------------------------------------#
library(spatstat)
#------------------------------------------------------------------#

# Read CSV into R
file.name <- "~/Desktop/TCGA/TCGA-GBM/Slide_Processing/TCGA-Processed/Matlab-Processed/csvs/batch03/TCGA-02-0339.csv"
MyObjectsData <- read.csv(file=file.name, header=TRUE, sep=",")

# Identify the x and y coordinates
x <- MyObjectsData$Centroid_1
y <- MyObjectsData$Centroid_2

# Create a point process object and display it
p <- ppp(x,y,c(1,max(x)),c(1,max(y)))
plot(p, pch=".")
# # Get the summary of the point process object
# summary(p)
ripley <- Kest(p)
plot(ripley, main = "", xlim=c(0, 30))
# #------------------------------------------------------------------#
# # Testing homogeneous Poisson
# #------------------------------------------------------------------#
# 
# M <- quadrat.test(p, nx=10, ny=10); M
# plot(M, add = TRUE, cex = 1)
# KS <- cdf.test(p, function(x, y) { x }); KS
# plot(KS)
# den <- density(p, sigma = bw.diggle(p))
# persp.im(den)
# contour.im(den)
# densities = solist(
#   `Diggle's edge correction` = density(subset(p), diggle = TRUE),
#   `No edge correction`       = density(subset(p), edge = FALSE)
# )
# plot(densities, equal.ribbon = TRUE, col = topo.colors)

#--------------------------------------------------------------------#
# We now fit Strauss process with 10 microns 
#--------------------------------------------------------------------#

# Fit the Strauss process and display
X <- ppm(p ~ polynom(x,y,2), Strauss(5))
X
# Computing and ploting the L-function of the dataset and choosing 
# the value r which maximises the discrepancy
plot(Lest(p), abs(iso - r) ~ r, main = "")
discrep <- function(r) {
  return(abs(as.function(Lest(p))(r) - r))
}
res <- optimise(discrep, interval = c(0, 10), maximum = TRUE)
print(res)

R <- res$maximum
X <- ppm(p ~ polynom(x,y,2), Strauss(R))
X
plot(fitin(X))
plot(Lest(p), main = "", xlim=c(0, 30))

rvals <- seq(1, 20, by = 0.1)
D <- expand.grid(r = rvals, sat=seq(1,3,by=1))
fitp <- profilepl(D, Geyer, p ~ polynom(x,y,2), aic=FALSE)
print(fitp)
plot(fitp)
summary(fitp)
X <- as.ppm(fitp)
X
fitp$fit$interaction$par[2]

# Run several simulations and see the validity of the model
diagnose.ppm(X, type = "Pearson", envelope = TRUE, nsim=40)
# Show the coefficients of the model
coef(X)
unlist(parameters(X))
# Plot the beta coefficients and a slice of the trend
plot(predict(X, type = "trend"))
plot(transect.im(predict(X, type = "trend")))
plot(transect.im(predict(X, type = "cif")))
#--------------------------------------------------------------------#
```
