## Supplementary material for "Gibbs Process Determines Survival and Reveals Contact-Inhibition Genes in Glioblastoma Multiforme": Codes and data: recursiveFitGeyer_AIC.R

```
#------------------------------------------------------------------#
# Load relevant libraries
#------------------------------------------------------------------#
library(spatstat)
#------------------------------------------------------------------#

# Read file names from the path into a vector
files <- list.files(path = "~/Desktop/TCGA/TCGA-GBM/Slide_Processing/TCGA-Processed/Matlab-Processed/csvs/batch09", full.names = TRUE)

# Start writing to an output file
# sink('~/Desktop/analysis-output_1.txt')

df <- data.frame(matrix(ncol = 13, nrow = 0))

for (file_index in 1:length(files)) {
  
  # Read each file and parse the data
  file.name <- files[file_index]
  
  # extract the name of the file 
  name.of.the.file <- basename(file.name)
  
  # print("--------------------------------------------------------------------")
  # print(file.name)
  # print("--------------------------------------------------------------------")

  MyObjectsData <- read.csv(file=file.name, header=TRUE, sep=",")
  
  # Identify the x and y coordinates
  x <- MyObjectsData$Centroid_1
  y <- MyObjectsData$Centroid_2
  
  # Create a point process object and display it
  p <- ppp(x,y,c(1,max(x)),c(1,max(y)))
  
  #--------------------------------------------------------------------#
  # We now fit Strauss process with 5 microns 
  #--------------------------------------------------------------------#
  
  # # Plot the L-function for each file
  # plot(Lest(p), abs(iso - r) ~ r, main = name.of.the.file)
  
  # Computing and ploting the L-function of the dataset and choosing
  # the value r which maximises the discrepancy

  discrep <- function(r) {
    return(abs(as.function(Lest(p))(r) - r))
  }
  res <- optimise(discrep, interval = c(0, 10), maximum = TRUE)

  # # Fit the Strauss process for the maximum L-function value 
  # strauss.parameter <- res$maximum
  # 
  # X <- ppm(p ~ polynom(x,y,2), Strauss(strauss.parameter))
  
  # Instead of the above fitting, we do a profilepl fit that computes 
  # pseudologlikelihood measures in the data frame defined by rvals
  rvals <- seq(1, 20, by = 0.1)
  D <- expand.grid(r = rvals, sat= seq(1, 3, by=1))
  # Fit the model for several data points
  fitp <- profilepl(D, Geyer, p ~ polynom(x,y,2), aic=FALSE)
  # Plot the AIC values (for checking correctness)
  plot(fitp)
  
  # Extract the best model
  X <- as.ppm(fitp)
  
  # Compute the interaction parameter gamma 
  gamma <- exp(X$coef[7])
  
  # Get the interaction distance
  interaction.distance <- fitp$fit$interaction$par[1]
  saturation.parameter <- fitp$fit$interaction$par[2]
  
  # We form the data frame and populate the computed coeffients 
  fit.coeff <- c(name.of.the.file, as.numeric(X$coef[1:7]), as.numeric(gamma), as.numeric(res), as.numeric(interaction.distance),
                 as.numeric(saturation.parameter))
  df <- rbind(df, as.data.frame(t(fit.coeff)))

}

# Assign columns to the data frame 
names(df) <-  c("Name","Intercept","x","y","I(x^2)","I(x * y)","I(y^2)","log(Interaction)","gamma","OptL","LObj","Interaction Distance",
                "Saturation Parameter")
df

# Write CSV in Desktop
write.csv(df, file = "~/Desktop/geyer_No_AIC_batch09.csv")

# sink()
```
