## Supplementary material for "Gibbs Process Determines Survival and Reveals Contact-Inhibition Genes in Glioblastoma Multiforme": Codes and data: segmentImage.m

```
%%
%% This program performs segmentation to the processed H&E images
%%

function segmentImage(image,cluster)

%%% Define full path to the file
filePath = '~/Desktop/TCGA/TCGA-GBM/Slide_Processing/TCGA-Processed/';
imageFile = strcat(filePath,image,'.png');

%%% Read the H&E image and display it
he = imread(imageFile);
% imshow(he), title('H&E image');

%%% Convert Image from RGB Color Space to L*a*b* Color Space
lab_he = rgb2lab(he);

%%% Classify the Colors in 'a*b*' Space Using K-Means Clustering
ab = lab_he(:,:,2:3);
ab = im2single(ab);
nColors = 3;

%%% Repeat the clustering 3 times to avoid local minima
pixel_labels = imsegkmeans(ab,nColors,'NumAttempts',3);

%%% Get the nucleus
mask2 = pixel_labels==cluster;
cluster2 = he .* uint8(mask2);
% figure; imshow(cluster2); title('Segmented nucleus');

%%% Convert the image into binary and display
bwImage = im2bw(cluster2,graythresh(cluster2));
% figure; imshow(bwImage)

%%% Erode the image to remove thinly connected nucleus
se = strel('line',4,4);
erodedBW = imerode(bwImage,se);
% figure; imshow(erodedBW)

%%% Remove small objects (less than 10 px)
rmSmallObjects = bwareaopen(erodedBW, 10);
% figure; imshow(rmSmallObjects)

%%% Fill the holes in the image
fillHoles = imfill(rmSmallObjects, 'holes');
% figure; imshow(fillHoles)

%%% Overlay the images
bwPerim = bwperim(fillHoles);
overlayedImage = imoverlay(he, bwPerim, [1 .3 .3]);
figure; imshow(overlayedImage)

%%% Counting the cells and other statistics
stats = regionprops('table',fillHoles,'Area', 'Centroid', ...
    'BoundingBox','MajorAxisLength','MinorAxisLength','Eccentricity', ...
    'Orientation','ConvexArea','FilledArea','EulerNumber', ...
    'EquivDiameter','Solidity','Extent','Perimeter','PerimeterOld');

%%% Check if the computed centroids are valid
% centroids = cat(1, stats.Centroid);
% hold on
% plot(centroids(:,1), centroids(:,2), 'yo')
% hold off

%%% Write the image and the table
writingFilePath = '~/Desktop/TCGA/TCGA-GBM/Slide_Processing/';

savedImageFilename = strcat(writingFilePath,image,'.png');
savedTableFilename = strcat(writingFilePath,image,'.csv');
savedBinaryFilename = strcat(writingFilePath,image,'_bw.png');

imwrite(fillHoles,savedBinaryFilename)
imwrite(overlayedImage,savedImageFilename)
writetable(stats,savedTableFilename)
```
